# Rehabilitation promotes the recovery of structural and functional features of healthy neuronal networks after stroke

**DOI:** 10.1101/582775

**Authors:** Anna Letizia Allegra Mascaro, Emilia Conti, Stefano Lai, Antonino Paolo Di Giovanna, Cristina Spalletti, Claudia Alia, Alessandro Panarese, Alessandro Scaglione, Leonardo Sacconi, Silvestro Micera, Matteo Caleo, Francesco Saverio Pavone

## Abstract

Rehabilitation is the most effective treatment for promoting the recovery of motor deficits after stroke. One of the most challenging experimental goals is to unambiguously link brain rewiring to motor recovery prompted by rehabilitative therapy. Here, we investigated which aspects of cortical remodeling are induced by rehabilitation by combining optical imaging and manipulation tools in a mouse model of stroke. We revealed that the stabilization of peri-infarct synaptic contacts fostered by rehabilitation goes along with increased vascular density induced by angiogenesis. Furthermore, we showed the progressive formation of a new motor representation in the peri-infarct area where temporal and spatial features of cortical activation recovered towards pre-stroke condition. In the same animals we observed the reinforcement of inter-hemispheric connectivity after rehabilitation. The present work provides the first evidences that rehabilitation promotes the combined recovery of structural and functional features distinctive of healthy neuronal networks.

## Introduction

Every year, several million stroke victims worldwide are impaired by long-term disability, imparting a large societal and personal burden in terms of lost productivity, lost independence and social withdrawal (Mozaffarian et al., 2015). When stroke affects the motor-associated cortices, the recovery of motor function is often partial, owing to the limited degree of spontaneous brain repair (Carmichael et al., 2017). Strategies that promote brain plasticity, like pharmacological treatments and physical training, can enhance neural rewiring and dramatically improve functional motor recovery. Pharmacological treatment both in the peri-infarct and contralesional areas has been shown to aid the restoration of function (for instance, see (Clarkson et al., 2010)). Another highly effective rehabilitative approach after stroke is physical training (Bütefisch, 2006; Jones and Adkins, 2015). The effectiveness of rehabilitative therapies can be maximized through a combination of treatments (Adkins-Muir and Jones, 2003; Dancause and Nudo, 2011; Hesse et al., 2007; Plautz et al., 2003). A few animal studies provided valuable insights into the anatomical adaptations in rodents and primates induced by coupled therapies (Fang et al., 2010; Lee et al., 2004; Plautz et al., 2003; Wahl et al., 2014). In a recent study on mice, Spalletti et al. took advantage of a rehabilitation paradigm that combines motor training and pharmacological inhibition of the contralesional primary motor cortex (M1) with Botulinum Neurotoxin E (BoNT/E) (Spalletti et al., 2017). BoNT/E injection reduced the excessive transcallosal inhibition exerted from the healthy to the stroke side (Spalletti et al., 2017). The combination of motor training and BoNT/E silencing was superior to either treatment alone in promoting recovery of motor skills in stroke mice. Importantly, this combined therapy led to motor improvements that generalized to multiple motor tasks (Spalletti et al., 2017). The induction of a generalized functional gain, i.e. the recovery of motor functions beyond the ones that are trained, is crucial when evaluating the efficacy of rehabilitative therapies.

In the last decade, fluorescence imaging provided valuable insights into spontaneous cortical plasticity after stroke. Structural plasticity after stroke has been evaluated by two-photon fluorescence (TPF) microscopy (Brown et al., 2007b; Johnston et al., 2013; Mostany et al., 2010; Sakadzic et al., 2015) showing that the extent of synaptic recovery varied along the distance from the infarct core (Sigler and Murphy, 2010) and that redistribution of blood vessel and dendrite orientation was confined to the peri-infarct area (Brown et al., 2007a). In parallel, Murphy and colleagues performed a number of acute optical imaging studies showing the persistence of cortical functional remapping over months after stroke (Brown et al., 2009; Harrison et al., 2013). The availability of a new generation of experimental tools, like optogenetics (Ayling et al., 2009; Lim et al., 2014) and genetically encoded functional indicators (Chen et al., 2013) provided new means of exploring neuronal rewiring.

To our knowledge, no investigation has yet described how rehabilitation after stroke affects cortical structural and functional plasticity *in vivo* and on the long-term scale. Many unanswered questions need to be addressed, like how rehabilitation molds neuronal structural plasticity and cortical functional maps. Moreover, it is still to be defined if neuronal and vascular plasticity act in concert to aid functional recovery.

In the present study, we dissected how rehabilitative treatment activates concurrent modalities of cortical plasticity in mice by using a combination of cutting-edge optical techniques. Our experiments revealed that a combination of physical training with pharmacological manipulation of the contralesional activity helps the preservation of dendritic architecture along with stabilization of spines in the peri-infarct region. We found in the same double-treated subjects the increase in the density of blood vessels primarily focused in the peri-infarct area. By longitudinally monitoring the calcium functional maps, we found that combined rehabilitation prompted the recovery of essential features of pre-stroke activation profiles, both in spatial and temporal terms. In the same animals presenting large-scale remapping of the injured hemisphere, we observed a significant enhancement of transcallosal functional connectivity after one month of rehabilitative therapy. Overall, our study provides an unprecedented view of complementary aspects of cortical anatomical and functional adaptation induced by rehabilitation after stroke.

## Results

### Experimental design

We used a photothrombotic model of focal stroke applied to the right M1 of GCaMP6f or GFPM mice (STROKE, TOXIN, ROBOT and REHAB groups). To restore the interhemispheric balance perturbed by stroke, we performed a focused inactivation of the forelimb motor cortex in the hemisphere contralateral to the ischemic lesion by means of intracortical injections of the synaptic blocker Botulinum Neurotoxin E (BoNT/E, TOXIN group). BoNT/E is a bacterial enzyme that enters synaptic terminals and reversibly blocks neurotransmission by cleaving SNAP-25, a main component of the SNARE complex (Caleo et al., 2007). We next tested the impact of physical rehabilitation by daily training the affected forelimb of stroke mice with controlled, repeatable and targeted exercises, guided by a robotic platform, namely the M-Platform (ROBOT group, (Spalletti et al., 2014)). Rehabilitation-associated training on the M-Platform consisted of repeated cycles of passively actuated contralesional forelimb extension followed by its active retraction triggered by an acoustic cue (Figure 1A). Finally, we tested the synergistic effect of combined therapy. The combined rehabilitation paradigm of the REHAB group consists in a combination of motor training on the M-Platform and pharmacological inactivation of the contralesional hemisphere via BoNT/E (Figure 1B).

**Figure 1:**
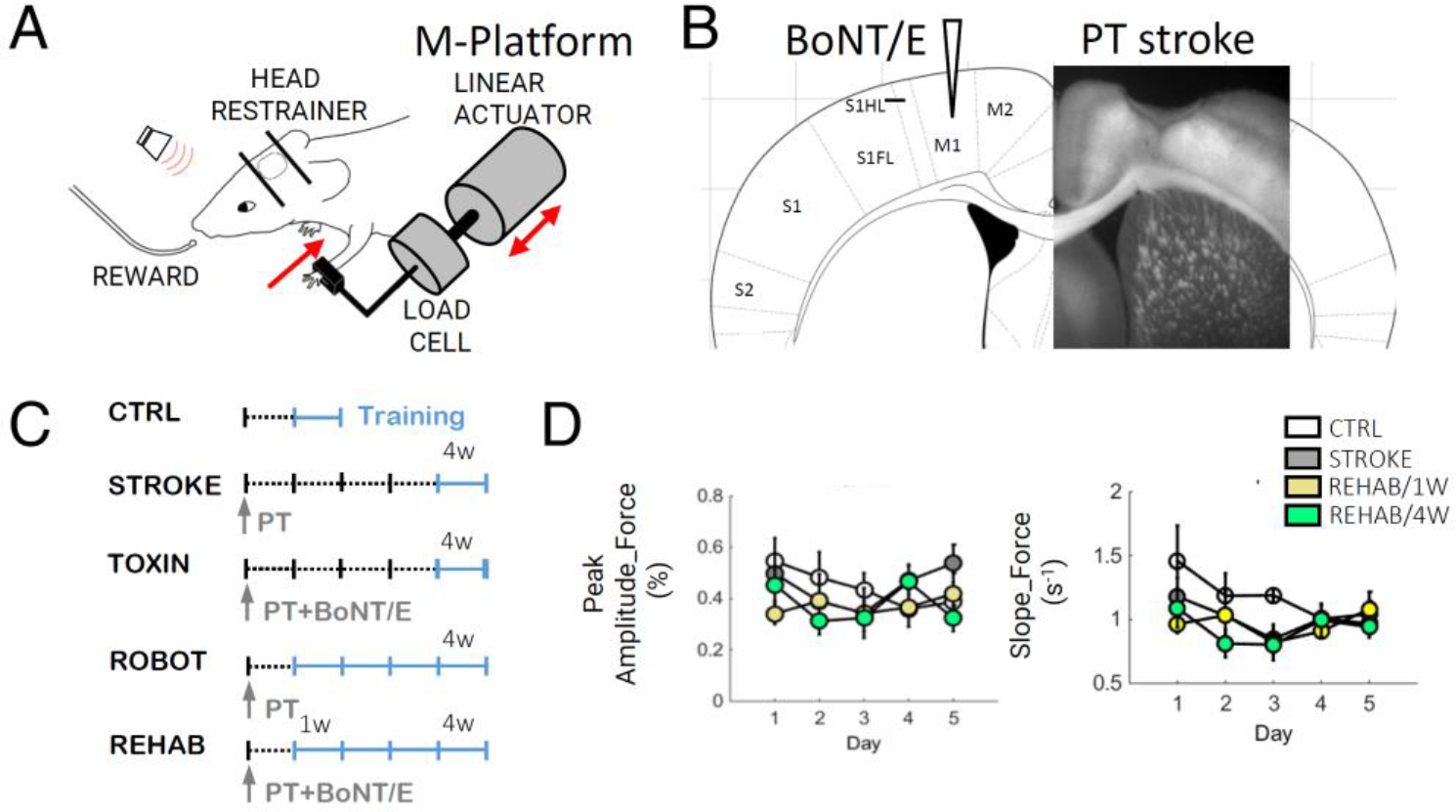
Experimental design. (A) Schematic representation of the M-Platform that was used for motor training. (B) Schematic of the experimental protocol, which combines the photothrombotic stroke in the primary motor cortex (M1) with a contralesional injection of BoNT/E into the homotopic cortex. (C) The experimental timeline for the CTRL, STROKE, TOXIN, ROBOT and REHAB groups in the awake imaging experiment. Light blue lines refer to training weeks. W = week; PT = photothrombosis. (D) Graphs showing the Peak amplitude (left) and Slope (right) of the force signals recorded in the pulling phase during training on the M-Platform over 5 days (4 weeks after injury for STROKE mice, 1 and 4 weeks after injury for REHAB mice, during the week of training for CTRL mice). There is no significant difference between the groups and over the 5 days.

ROBOT and REHAB mice were trained for 4 weeks starting 5 days after injury (Figure 1C), in line with the overall consensus that the initiation of rehabilitative training 5 or more days after stroke is mostly beneficial and has no adverse effects (Krakauer et al., 2012). The motor task was rapidly learned and easily performed. Indeed, the amplitude and slope of the force peaks exerted during the voluntary forelimb-pulling task were not significantly different across groups, neither within a week nor across weeks of training (Figure 1D).

### Combined rehabilitation preserves pyramidal neurons architecture and promotes the stabilization of synaptic contacts in the peri-infarct area

In the present study we assessed how different rehabilitative approaches prompt large- and small-scale cortical plasticity. In particular, we wondered if the improved performance in generalized motor tasks induced exclusively by the combined treatment (Spalletti et al., 2017) resulted in distinct features of cortical reorganization. We investigated synaptic dynamics, vascular network remodeling and remapping of motor representation.

First, we explored structural rewiring of dendrites and spines. Heightened spontaneous structural remodeling was observed in mice at both presynaptic (axonal fibers and terminals) and postsynaptic (dendrites and dendritic spines) sites in the peri-infarct cortex in the weeks following stroke (Brown et al., 2010; Carmichael et al., 2001; Dancause et al., 2005; Hsu and Jones, 2006; Ueno et al., 2012). In this set of *in vivo* experiments, we performed TPF microscopy of pyramidal apical dendrites and axons (0-100 μm deep from the pial surface) with increasing distances (up to 4 mm) from stroke core in the cortex of GFPM mice (Figure S1A). Orientation of dendrites and axons was evaluated on the 4^th^ week after performing the lesion (STROKE, TOXIN, ROBOT and REHAB mice).

A strong orientation gradient of dendrites induced by the collapse of dead tissue in the ischemic core was evident in the STROKE group (Figure 2A, right panels and Figure 2B). Reorientation was not detectable at distal regions from the ischemic core (>1 mm; Figure S1B and S1C). A strong alignment of neurites in the peri-infarct region was observed in stroke mice treated with BoNT/E (TOXIN group, Figure 2B). Four weeks of robotic training (ROBOT group) were not sufficient to prevent the formation of this structural abnormality. In contrast, reorientation was not visible in REHAB animals, nor in proximal (Figure 2B) or distal (Figure S1C) regions. Indeed, the randomness in dendrite orientation in REHAB mice resembled the one observed in healthy (CTRL) subjects (Figure 2B).

**Figure 2:**
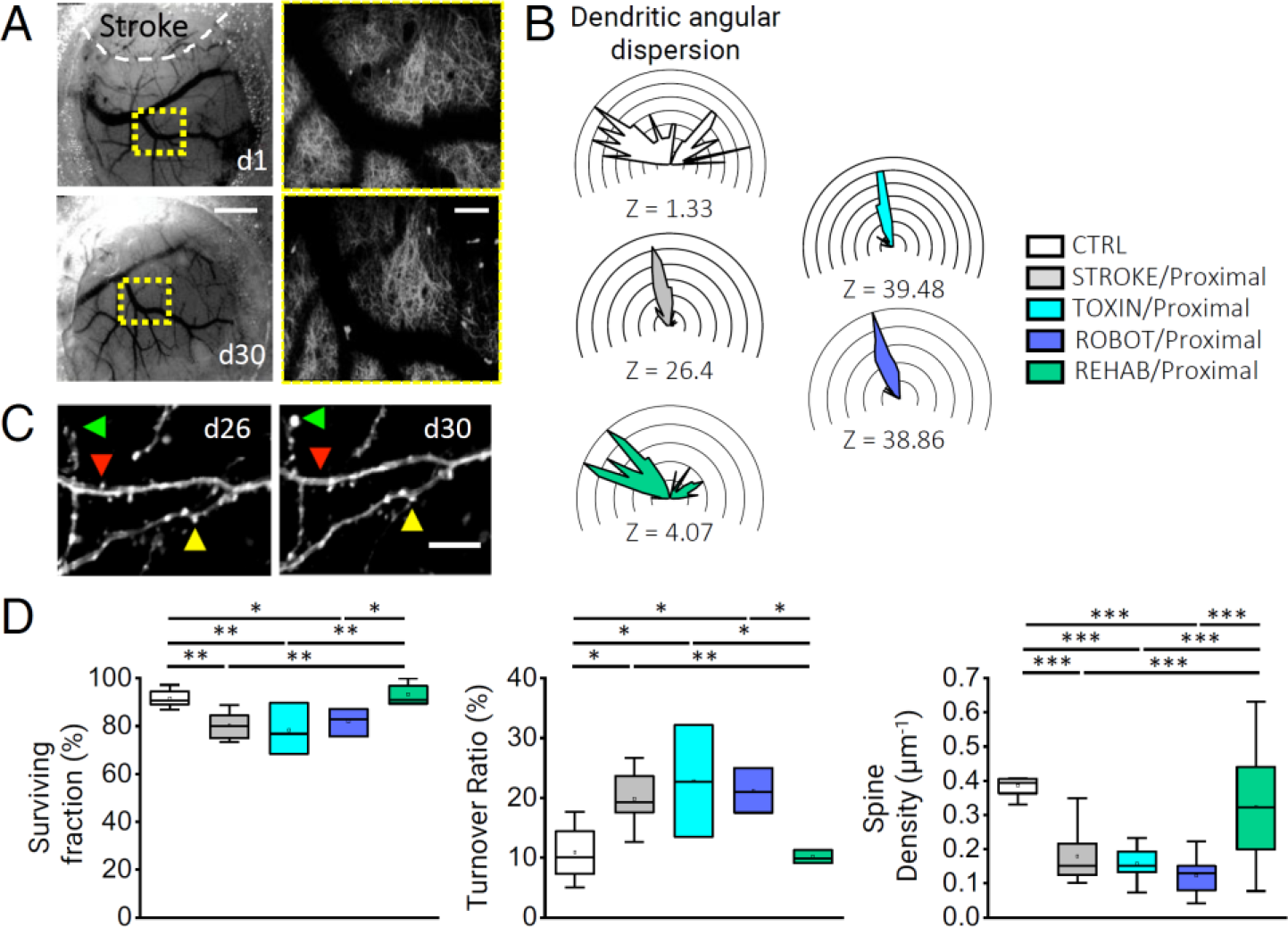
Combined rehabilitation affects dendritic orientation and stabilizes spine turnover. (A) Brightfield images showing cranial window 1 day and 30 days after stroke (STROKE group). The rostral shift of cortical tissue due to the shrinkage of the stroke core is highlighted by the displacement of a reference point (i.e. blood vessel branching point) framed by the dotted yellow square. Scale bar, 1 mm. Panels on the right show the stitched two-photon images (4×4 Maximum Intensity Projections, MIPs of 130×130×50 µm^3^) acquired within the region framed by the respective yellow squares in the left panels. Scale bar, 100 μm. (B) Polar plots showing angular distribution of dendrites in peri-infarct area. Plots are oriented as in Figure 2A, where lesioned area is toward the uppermost center of the plot (for all panels, Nmice_CTRL_ = 5, Nmice_STROKE_ = 6, Nmice_REHAB_ = 5). Z scores, calculated by Rayleigh Test for circular statistics, are reported for each experimental class at the bottom of the polar plot. (C) MIPs of two-photon stacks (z depth 8 μm) of dendritic branches at 26 and 30 days after stroke (STROKE group). Arrowheads point to a newly formed (green), lost (red), and stable (yellow) spine. Scale bar, 5 μm. (D) Box and whiskers plot showing surviving fraction (SF) ± SEM (*left*; SF_CTRL_ = 92 ± 2%; SF_STROKE_ = 80 ± 2%; SF_TOXIN_ = 78 ± 6%; SF_ROBOT_ = 81 ± 3%; SF_REHAB_ = 93 ± 2%; one-way ANOVA with post hoc Fisher test, *P_CTRL/STROKE_* = 0.006, *P_REHAB/STROKE_* = 0.002, *P_CTRL/TOXIN_* = 0.008, *P_REHAB /TOXIN_* = 0.003, *P_CTRL/ROBOT_* = 0.04, *P_REHAB /ROBOT_* = 0.02), turnover ratio (TOR) ± SEM (*middle*; TOR_CTRL_ = 11 ± 2%; TOR_STROKE_ = 20 ± 2%; TOR_TOXIN_ = 23 ± 5%; TOR_ROBOT_= 21 ± 2%; TOR_REHAB_ = 10 ± 3%; one-way ANOVA with post hoc Fisher test, *P_CTRL/STROKE_* = 0.02, *P_REHAB /STROKE_* = 0.01, *P_CTRL/TOXIN_* = 0.01, *P_REHAB /TOXIN_* = 0.008, *P_CTRL/ROBOT_* = 0.03, *P REHAB/ROBOT* = 0.02) and spine density (SD) ± SEM (*right panel*; SD_CTRL_ = 0.39 ± 0.01 µm^-1^; SD_STROKE_ = 0.18 ± 0.02 µm^-1^; SD_TOXIN_ = 0.16 ± 0.01 µm^-1^; SD_ROBOT_ = 0.12 ± 0.02 µm^-1^; SD_REHAB_ = 0.32 ± 0.05 µm^-1^; one-way ANOVA with post hoc Bonferroni test *P_CTRL/STROKE_* = 4.1*10^-6^, *P_REHAB/STROKE_* = 5.5*10^-4^, *P_CTRL/TOXIN_* = 1.0*10^-6^, *P REHAB/TOXIN* = 1.2*10^-4^, *P_CTRL/ROBOT_* = 4.3*10^-6^, *P_REHAB/ROBOT_* = 2.6*10^-4^) in the peri-infarct area. See also Figure S1.

Alterations induced by ischemic damage on neural circuitry are known to affect dendrites as well as synaptic contacts, producing a large increase of spine turnover in peri-infarct area of spontaneously recovering mice (Brown et al., 2007a; Mostany et al., 2010). We next asked how different rehabilitative approaches modified synaptic turnover. We thus monitored the appearance and disappearance of apical dendritic spines (Figure 2C) by performing a frame-by-frame comparison of spines in the mosaic we acquired along the rostro-caudal axis with TPF microscopy. We first evaluated the peri-infarct plasticity of pyramidal neurons in spontaneously recovering mice (STROKE) and verified that this focal lesion triggers an increased instability of synaptic contacts. Surprisingly, the single treatments by inhibition of the contralesional hemisphere or physical therapy alone were not able to reinstate physiological levels of spine plasticity (Figure 2D).

On the contrary, spines in the peri-infarct region of REHAB animals exhibited increased synaptic stability (Figure 2D, left panel) and a lower turnover (Figure 2D, middle panel) than STROKE mice, that were comparable to healthy CTRL values. The combined rehabilitative treatment also resulted in higher synaptic densities thereby recapturing an important feature of pre-stroke conditions (Figure 2D, right panel). The stabilization of synaptic contacts induced by rehabilitation was stronger at the proximal level and weakened with increasing distance from ischemic core (Figure S1D, upper and middle panels). Interestingly, the change in density of spines extended beyond peri-infarct region (Figure S1D, lower panel).

In brief, longitudinal imaging of cortical neurons revealed that the combined rehabilitative therapy was necessary to preserve the organization of dendritic arbors in peri-infarct cortex and to restore dendritic spine plasticity.

### Combined rehabilitative treatment stimulates angiogenesis in the peri-infarct area

Since stroke profoundly altered the spatial distribution of superficial blood vessels (Figure 2A, left panels), we wondered if and how rehabilitation altered vascular milieu in the ipsilesional cortex. To test this, we longitudinally observed mouse cortices of all groups under both cranial window and thinned skull preparations for one month after stroke. We found that the injured area progressively shrunk owing to the collapse of dead tissue in STROKE and REHAB mice (Figure 2A, 3A and S2). In addition, a very bright area appeared in the peri-infarct region of both groups of GCaMP6 mice, possibly representing an excitotoxic response associated with calcium dysregulation (Ankarcrona et al., 1995) elicited by the photothrombotic stroke (see the left panel of Figure 3A; 13 out of 14 STROKE and REHAB mice). The enhanced brightness gradually diminished and disappeared after the acute period (6-19 days after injury in STROKE and REHAB mice; see examples in Figure 3A, right panel and Figure S2), and was accompanied by a large shrinkage of the necrotic tissue. The consequent displacement of the peri-infarct area was associated with a substantial remodeling of blood vessels (white arrowheads in Figure 3A), in agreement with previous studies (Brown et al., 2007a).

**Figure 3:**
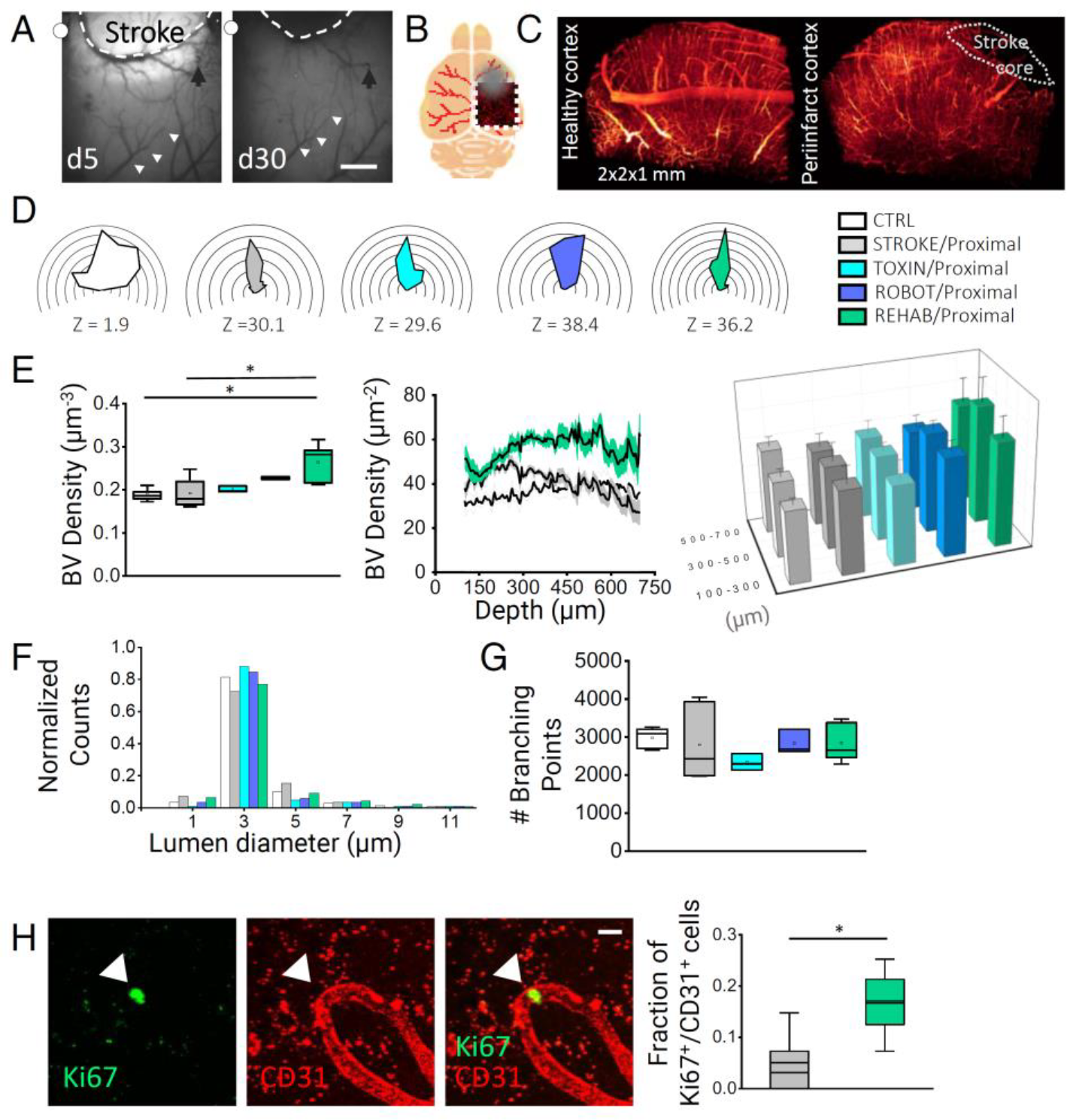
Combined rehabilitation promotes angiogenesis in the peri-infarct area. (A) Brightfield images showing cortical vasculature after stroke under a thinned skull preparation. Dotted line emphasizes the profile of a large blood vessel shifting toward the stroke core from 5 days to 30 days after stroke; a similar shift is highlighted by the white arrowheads on a blood vessel distal to the injury site. Black arrow points to an internal reference on the image. White dots indicate bregma. Scale bar, 1 mm. (B) Schematic representation of fixed brain of a GFPM mouse, in which vasculature is labeled with TRITC (red lines). Dotted square highlights the imaged area; gray circle labels stroke location. (C) 3D meso-scale reconstruction of vasculature in the healthy contralesional cortex (left) and in the peri-infarct area (right) from TDE-cleared cortical slices. White dotted region highlights the absence of blood vessels in the infarct core. (D) Polar plots show the distribution of blood vessel orientation in regions proximal to the core for each experimental group measured after the last motor training session (Nmice_CTRL_ = 5; Nmice_STROKE_ = 6; Nmice_TOXIN_ = 3; Nmice_ROBOT_ = 3; Nmice_REHAB_ = 5). Plots are oriented as in Figure 3A; the lesioned area is toward the uppermost center of the plot. Z scores, calculated by the Rayleigh Test for circular statistics, are reported for each experimental class below the plot. (E) Blood vessel (BV) density analysis. *(Left)* The box and whiskers plot shows the BV density (average ± SEM) in regions proximal to the stroke core in each experimental group (BV Density_CTRL_ = 0.19 ± 0.01 µm^-3^; BV Density_STROKE_ Proximal = 0.19 ± 0.03; BV Density_TOXIN_ Proximal = 0.20 ± 0.01; BV Density_*ROBOT Proximal*_= 0.23 ± 0.01; BV Density_*REHAB Proximal*_ = 0.26 ± 0.05; one-way ANOVA with post hoc Bonferroni test, *P_REHAB Proximal/STROKE Proximal_* = 0.018, *P*_*REHAB Proximal/CTRL*_ = 0.017). *(Middle)* Traces indicate BV density with respect to brain cortex depth in each experimental group. Shadows represent SEM. *(Right)* 3D graph comparing the average BV density grouped by cortical depth (100-300 µm from stroke core: BV Density_CTRL_ = 0.18 ± 0.02; BV Density_STROKE Proximal_ = 0.20 ± 0.04; BV Density_TOXIN Proximal_ = 0.19 ± 0.05; BV Density_ROBOT Proximal_ = 0.24 ± 0.03; BV Density_REHAB Proximal_ = 0.26 ± 0.05; 300-500 µm from stroke core: BV Density_CTRL_ = 0.18 ± 0.02; BV Density_STROKE Proximal_ = 0.20 ± 0.04; BV Density_TOXIN Proximal_ = 0.21 ± 0.02; BV Density_ROBOT Proximal_ = 0.24 ± 0.02; BV Density_REHAB Proximal_ = 0.29 ± 0.06; 500-700 µm from stroke core: BV Density_CTRL_ = 0.20 ± 0.02; BV Density_STROKE Proximal_ = 0.18 ± 0.03; BV Density_TOXIN Proximal_ = 0.20 ± 0.03; BV Density_ROBOT Proximal_ = 0.20 ± 0.03; BV Density_REHAB Proximal_ = 0.24 ± 0.06; 300-500 µm from the stroke core: *P*_*REHAB Proximal /STROKE Proximal*_ = 0.024, *P*_*REHAB Proximal /CTRL*_ = 0.013; one-way ANOVA with post hoc Bonferroni test). (F) Histogram showing the distribution of lumen diameter of BV for CTRL mice and in the proximal region of STROKE, TOXIN, ROBOT and REHAB mice. (G) The box and whiskers plot shows the quantification of the number of branching points in proximal regions for each experimental group (Number of Branching Points (BP): BP_CTRL_ = 2979 ± 126; BP_STROKE_ = 2797 ± 390; BP_TOXIN_ = 2332 ± 126; BP_ROBOT_ = 2833 ± 185; BP_REHAB_ = 2852 ± 243). (H) Immunohistochemical analysis on brain slices with double Ki67 and CD31 labeling. On the left, the images show one example of colocalization of a Ki67^+^ cell within a CD31^+^ endothelial cell within a blood vessel in the peri-infarct cortex. The white arrowhead points at the location of the Ki67^+^ cell in the three panels. On the right, box and whiskers plot reports the fraction of double stained Ki67^+^/CD31^+^ cells in STROKE (N=5) vs REHAB (N=5) mice (Fraction of Ki67^+^/CD31^+^cells STROKE= 0.05 ± 0.03; REHAB= 0.17 ± 0.03; one-tailed P-value = 0,012). Scale bar, 10 µm. See also Figure S2, S3 and Video S1.

We quantified the structural reshaping of cortical vasculature with high resolution by performing 3D reconstructions with TPF microscopy. Specifically, after the last motor training session, we stained the brain vasculature of GFPM mice with a vessel-filling fluorescent dye, albumin-fluorescein isothiocyanate (FITC, Figure 3B). By imaging Thiodiethanol (TDE) cleared cortical slices (see Methods), we obtained high-resolution 3D maps of blood vessels of the injured, right hemisphere (Figure 3C, Video S1). The reorganization of cortical tissue (Figure 2A) that led to a considerable alignment of dendrites towards the stroke core produced an analogous re-orientation of blood vessels in peri-infarct area of STROKE mice (Figure 3D). Interestingly, neither the single treatment (TOXIN and ROBOT mice) nor combined rehabilitation (REHAB) could rescue random vascular orientation of pre-stroke conditions in the peri-infarct (proximal, <500 μm) region (Figure 3D). A less pronounced orientation was visible in all groups at distal regions (1000 to 1500 μm from core, Figure S3A).

We further guessed if rehabilitation triggered angiogenesis in the ipsilesional cortex. We found that the single treatments (TOXIN and ROBOT groups) did not have any significant impact on blood vessel numbers (Figure 3E, left panel). Conversely, the density of blood vessels near the infarct site was significantly higher in REHAB mice compared to both STROKE and CTRL groups.

We then evaluated the vascular density over the entire cortical depth to define layer-specific contributions to the overall increase in blood vessel density. Blood vessel distribution was quite uniform in CTRL mice. Increases in density were localized mainly to the middle layers of the cortex (300-500 μm deep) in REHAB group (Figure 3E, middle and right panels). We found that the augmented vascular density was mainly observed in regions near the stroke core (Figure 3E, right panel), whereas distal regions did not show any significant increase (Figure S3B).

We then asked if the increased density was due to an increase in average diameter of blood vessels. Our results show that there is no significant difference in distribution of lumen sizes of cortical blood vessels for the five experimental groups (Figure 3F), ruling out possible influences of vasodilation or vasoconstriction in the evaluation of density. Alternatively, the increased density could be due to ramification of existing vessels, leading to an increase in the complexity of the vascular maps. Nevertheless, the evaluation of the number of branching points showed no significant differences between the spontaneously recovering mice (STROKE) and all the treatments (TOXIN, ROBOT, REHAB) nor in proximal (Figure 3G) or distal (Figure S3C) regions, thus ruling out this hypothesis.

Finally, we asked if rehabilitation fostered an increase in endothelial proliferation that eventually led to augmented vascular density. We thus compared a new group of REHAB mice after 10 sessions of daily robotic training with spontaneously recovering mice (STROKE group) 16 days after stroke. To evaluate endothelial proliferation, we performed ex vivo double labeling on brain slices with CD31 and Ki67 (see Supplementary Material section). Figure 3H shows that the number of double stained cells is higher in the peri-infarct region of the cortex of REHAB compared to STROKE mice. The augmented endothelial cell proliferation in combination with the increased density of blood vessels (higher than healthy CTRL mice) and the non-significant change in lumen diameter in REHAB mice strongly suggest that the combined treatment promotes angiogenesis in the peri-infarct area.

Taken together, these results on cortical vasculature indicate that the combination of pharmacological and physical therapy triggers substantial blood vessel growth via the promotion of endothelial cell proliferation in the peri-infarct region.

In conclusion, our results suggest that generalized functional recovery is associated with a pro-angiogenic effect on the vasculature of the peri-infarct area induced by the combined rehabilitative treatment.

### Combined rehabilitation treatment restores cortical activity patterns disrupted by stroke

We next assessed if combined rehabilitation modulated cortical motor representation activated by voluntary movement of the injured forelimb. We implemented an integrated system for simultaneous imaging of the calcium indicator GCaMP6f over the injured hemisphere and recording of forces applied by the contralesional forelimb during the training sessions on the M-Platform (Figure 4A-C). Calcium imaging was used as a measure of cortical activity in the brains of GCaMP6f mice. We focused on the analysis of calcium response activated in the same time-window of the voluntary retraction movement (see example in Figure 1C, right panel). The imaging area overlapped with the region visualized in GFPM mice (Figure 4B). Wide-field calcium imaging showed that a small area located in the motor-sensory region reproducibly lit up in CTRL mice during the forelimb retraction movement on the M-platform (examples of cortical activation are reported in Figure 4D and Figure S4A). On the contrary, a large area covering most of the cortical surface of the injured hemisphere was activated while performing the task in non-treated (STROKE) mice one month after stroke. Remarkably, calcium activation in REHAB mice was similar to healthy controls (CTRL) in terms of extension, location, timing and amplitude (Figure 4D-H, Figure S4A-D).

**Figure 4:**
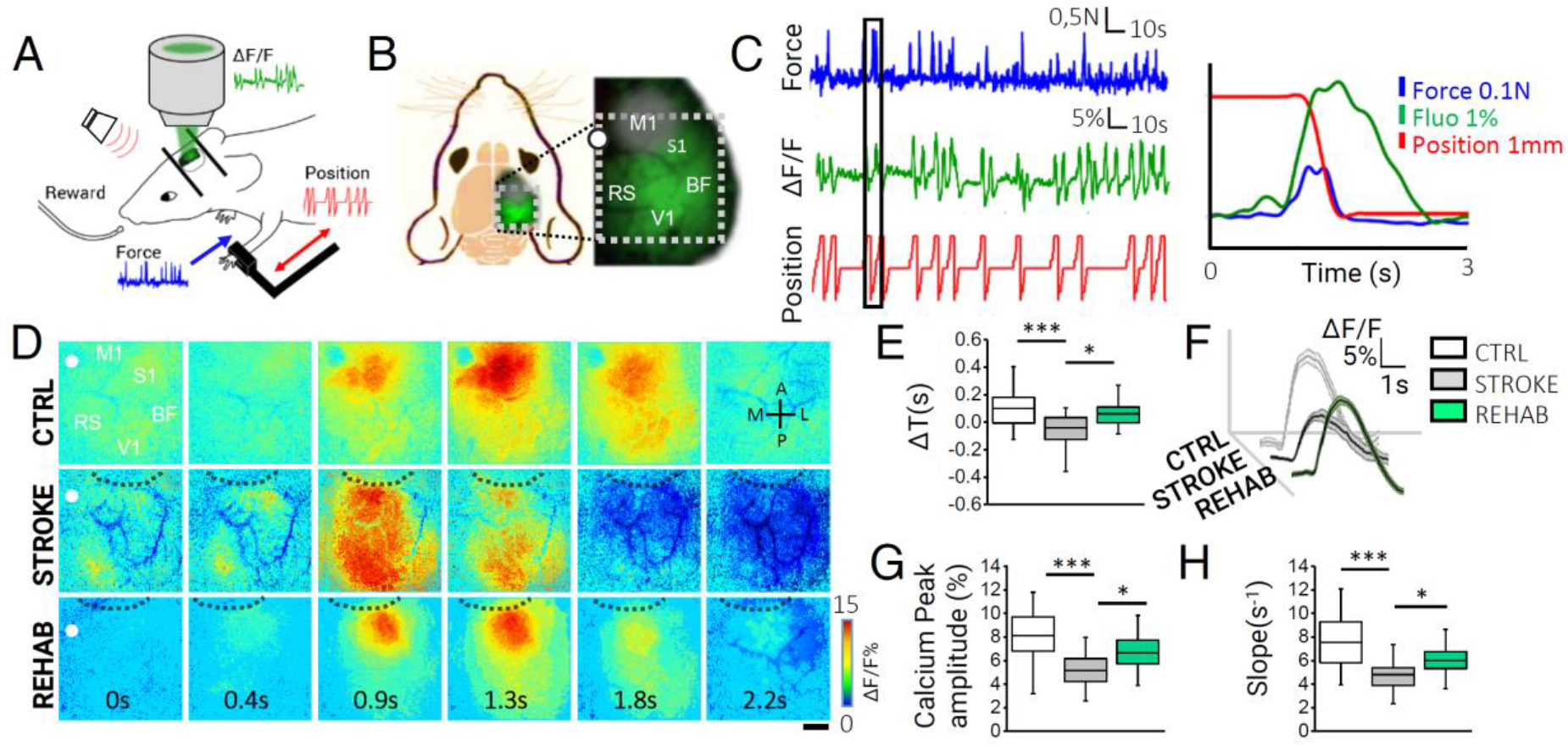
Combined rehabilitation counteracts cortical dedifferentiation and restores activation profiles in the post stroke, peri-infarct region. (A) Schematic representation of M-Platform: examples of simultaneously recorded force (blue), position (red) and ΔF/F traces (green) are reported. (B) Schematic representation of field of view (i.e. area within the dotted white square) used for wide-field calcium imaging in GCaMP6f mice. The gray circle on M1 region highlights the location and approximate extent of the lesion. M1, primary motor area; S1, primary somatosensory area; BF, barrel field; V1, primary visual cortex; RS, retrosplenial cortex. White dot indicates the bregma. (C) Example of force trace (blue), fluorescence trace (green) and handle position (red). The graph on the right shows an overlap of simultaneously recorded traces corresponding to the black box on the left. (D) Image sequence of cortical activation as assessed by calcium imaging during pulling of the handle by the contralateral forelimb, from 0.4 s before to 1.8 s after the onset of the force peak. Each row shows a representative sequence from a single animal of each group. Black dashed lines define the lesion borders. Scale bar, 1mm. (E) Delays in cortical activation in caudal regions in response to contralateral forelimb retraction are shown for the 3 groups (Nmice_CTRL_ = 4, Nmice_STROKE_ = 6, Nmice_REHAB_ = 6; ΔT_CTRL_ = 0.10 ± 0.03 s, ΔT_STROKE_ = −0.04 ± 0.02 s, ΔT_REHAB_ = 0.06 ± 0.02 s; Kruskal-Wallis One Way followed by Tukey’s test: *** *P* = 0.006, * *P* = 0.016). (F) Average calcium traces recorded in the region of maximal activation, ROI_g_ (see Methods), during contralateral forelimb retraction for the experimental groups; the shadows indicate SEM values. (G) Graph shows the maximum of fluorescence peaks from the same calcium traces as in (F) (Peak amplitude_CTRL_ = 8.1 ± 0.5 %; Peak amplitude_STROKE_ = 5.2 ± 0.3%; Peak amplitude_REHAB_ = 6.6 ± 0.3%; one-way ANOVA followed by Tukey’s test: *** *P* = 0.000005, * *P* =0.017). (H) Graph shows the slope (average ± SEM) of the fluorescence in the rising phase of the trace (Slope_CTRL_ = 7.5 ± 0.5 s^-1^; Slope_STROKE_ = 4.8 ± 0.2 s^-1^, Slope_REHAB_ = 6.0 ± 0.2 s^-1^; Kruskal-Wallis One Way followed by Tukey’s test: *\*\*\* P* = 0.00004, * *P* = 0.015). See also Figure S4.

We analyzed the extension and location of the motor representation by overlapping the movement-triggered activation maps obtained on every day of the training week (*ROIg*, see Figure S4B-C and Supplemental Procedures). Stroke expanded the motor representation from M1 toward more caudal regions not specifically associated with motor control (STROKE group in Figure S4C, 2^nd^ panel). Interestingly, the extension of motor representation in REHAB mice was consistently reduced (Figure S4C 3^rd^ panel and Figure S4D). In terms of location, the motor representation of REHAB mice was centered on the motor-associated region in the peri-infarct area (Figure S4C 3^rd^ panel and Figure S4D). The localization and spread of motor maps changed progressively along the weeks of training (Figure S4F). In most cases (5 out of 6 REHAB mice) we found a higher correlation (i.e. augmented functional connectivity) in the activity of spared motor-associated areas (Figure S4E) at the end of the training period (REHAB/4W) compared to the beginning (REHAB/1W). We hypothesized that focalized versus spread motor representations could be associated with different patterns of propagation of calcium activity. To quantify the concurrent recruitment of motor-associated and other functional areas, we analyzed the temporal profile of movement-triggered calcium transients over the injured hemisphere. We assumed that an extended activation (as in STROKE mice) implied a synchronicity in the activation of the rostral peri-infarct and the caudal areas. Indeed, we found that the delay between the maximum calcium peak of the rostral and caudal regions was slightly negative in STROKE mice, indicating that the activation of the caudal region somewhat preceded the activation of the rostral (peri-infarct) one (Figure 4E). Conversely, the delay was positive in CTRL and REHAB mice. Longitudinal imaging showed that the temporal pattern of rostro-caudal calcium activation in REHAB mice was gradually recovered towards pre-stroke conditions during 4 weeks of training (Figure S4G, left panel).

To sum up, the combined rehabilitative treatment focalized the motor representation to the peri-infarct region. The motor representation of REHAB mice closely resembled the one observed in CTRL animals both spatially and temporally.

We further analyzed calcium transients in peri-infarct area during contralateral forelimb retraction in CTRL, STROKE, and REHAB mice. At a glance, the average traces in Figure 4F show that stroke (in STROKE mice) and the combined treatment (REHAB mice) modified several features of the calcium transients. While a significant reduction in the amplitude of calcium transients during forelimb retraction was evident one month after stroke in the STROKE group, REHAB mice partially recovered to pre-stroke conditions (Figure 4G). In detail, the combined treatment led to a progressive increase in the amplitude of the calcium response over the training period (Figure S4G, middle panel). Moreover, REHAB animals showed progressively steeper slope of calcium activation along the weeks of training (Figure S4G, right panel). As compared to STROKE animals, a faster rise of calcium transient in the peri-infarct region was observed in REHAB mice at the end of the training period (Figure 4H).

In brief, our results showed that combined rehabilitative treatment promoted the formation of a new motor representation in the peri-infarct area where temporal and spatial features of cortical activation recovered towards pre-stroke condition.

### Combined rehabilitation improves inter-hemispheric functional connectivity

We hypothesized that the combined treatment alters the functional connectivity of the new motor representation with the contralesional motor cortex. Several studies in mice and humans have shown that interhemispheric M1 connectivity is reduced after stroke (Bauer et al., 2014; van Meer et al., 2010a; van Meer et al., 2010b). Therefore, it is hypothesized that increased interhemispheric connectivity positively correlates with the recovery of motor performances in the subacute stage after stroke in humans (Carter et al., 2010). Based on this hypothesis, we tested whether transcallosal projections were altered by combined rehabilitation by using an all-optical approach that combined optogenetic activation of the intact M1 with calcium imaging on the injured hemisphere. In these experiments, the intact (left) M1 of GCaMP6f mice was injected with AAV9-CaMKII-ChR2-mCherry to induce the expression of ChR2 in pyramidal neurons (Figure 5A).

**Figure 5:**
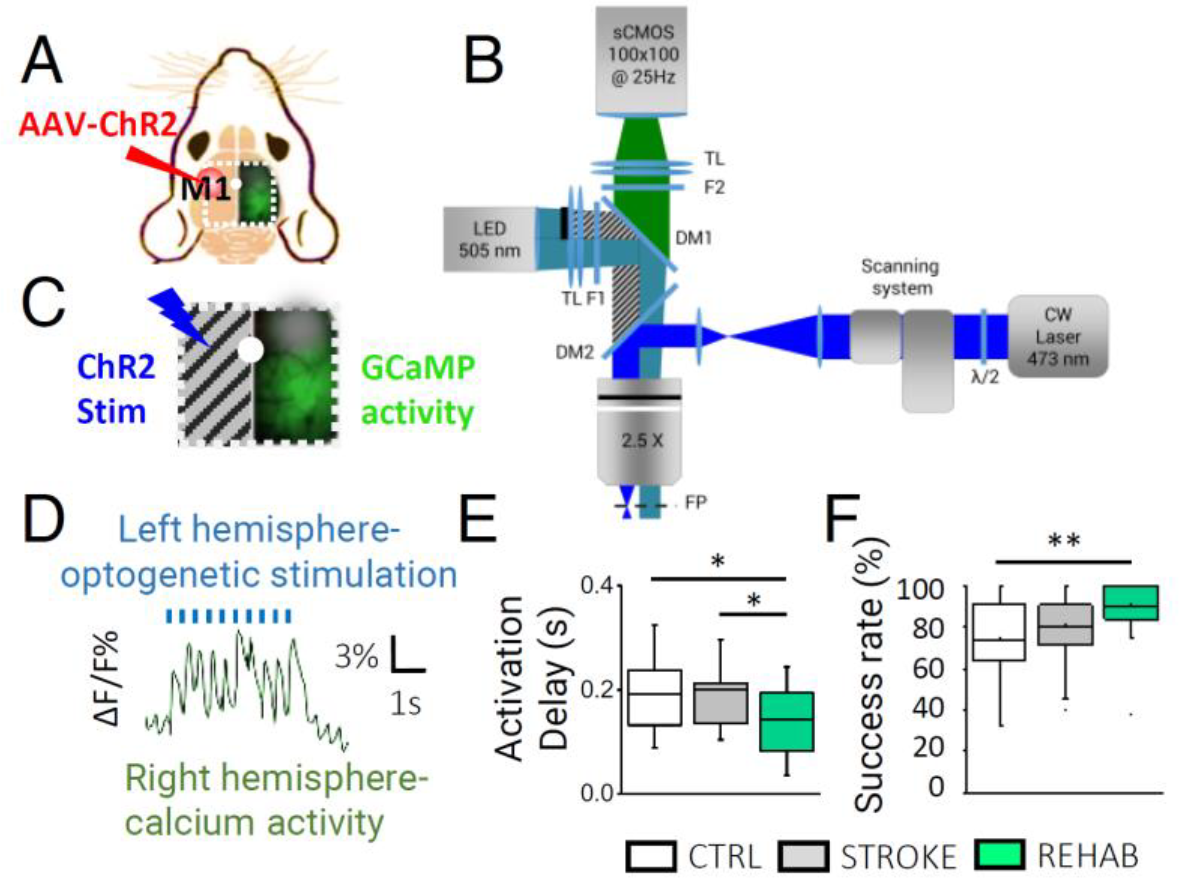
Combined rehabilitation strengthens interhemispheric connectivity after stroke. (A) Schematic representation of field of view (i.e. area within the white dotted square) for all-optical investigation of inter-hemispheric functional connectivity in GCaMP6f mice. The red circle in the left hemisphere indicates M1 injected with AAV-ChR2-mCherry. White dot indicates bregma. (B) Schematic representation of wide-field microscope. (C) Panel shows simultaneous ChR2 laser stimulation on the left hemisphere (not illuminated by LED) and the detection of the evoked GCaMP6f fluorescence on the right side. (D) Trace of optogenetically elicited calcium activity in the right cortex. (E) Box and whiskers plot showing the delay between the onset of left hemisphere laser stimulation and the peak of the calcium responses in the right hemisphere (average ± SEM; Nmice_CTRL_ = 4, Nmice_STROKE_ = 3, Nmice_REHAB_ = 3; Activation Delay_CTRL_ = 0.19 ± 0.02 s, Activation Delay_STROKE_ = 0.20 ± 0.02 s, Activation Delay_REHAB_ = 0.14 ± 0.01 s; one-way ANOVA post hoc Fisher, *P* <0.05 for all comparisons;). (F) Success rate (SR) of laser stimulation, calculated as number of times laser stimulation successfully triggered contralateral activation over total number of stimulation trials (average ± SEM, same Nmice as in (E); SR_CTRL_ = 74 ± 4, SR_STROKE_ = 81 ± 3, SR_REHAB_= 90± 3; ANOVA with post hoc Bonferroni test: *P* < 0.01). See also Figure S5 and Video S2.

One month after stroke, we optogenetically stimulated the intact M1. To avoid ChR2 activation while exciting GCaMP6f fluorescence, we partially occluded the 505 nm LED in our custom-made wide-field microscope (Figure 5B-C; see also (Conti et al., 2019)). Optogenetic stimulation was achieved with a second excitation path in which a 470 nm laser was focused on the AAV-transfected region via acousto-optic deflectors (AODs) (Crocini et al., 2016).

Laser stimulation of the contralesional M1 reproducibly triggered the activation of calcium transients in the injured hemisphere (Figure 5D, Video S2), mainly spreading from the homotopic M1 of CTRL mice, or in the peri-infarct area of STROKE and REHAB animals (Figure S5). Cortical activity then propagated to functionally connected regions that were located either anterior or posterior to the primary source of activation in all 3 groups of animals (Video S2). By quantifying the delay between the start of the optogenetic stimulation on the left hemisphere and the peak of calcium activity in the right hemisphere, we found no significant difference between the STROKE and CTRL mice (Figure 5E). In contrast, this delay was significantly reduced in REHAB mice as compared to both CTRL and STROKE mice (Figure 5E). In addition, the success rate of contralateral activation in response to optogenetic stimulation was higher in REHAB mice than in CTRL and STROKE animals (Figure 5F). Therefore, this series of all-optical experiments suggests that combined rehabilitation strengthens the functional connectivity between the spared motor cortex and the perilesional cortex one month after stroke.

## Discussion

In the present study we showed that rehabilitative treatment combining physical training and BoNT/E inhibition of contralesional hemisphere stabilized perilesional synaptic contacts, restored features of cortical activation typical of pre-stroke conditions and promoted the formation of an enriched vascular environment to feed the neuronal population that has been recruited to perform motor control. These results offer novel insights on cortical plasticity induced by rehabilitation *in vivo*. To the best of our knowledge, this study provides first-time evidence on the correlation between rehabilitation-induced neuronal and vascular reshaping in the mouse cortex after stroke.

The combined rehabilitation paradigm used here was recently characterized in terms of behavioral assessment of motor recovery (Spalletti et al., 2017). The authors previously developed the M-Platform, a mechatronic device for mouse forelimb training (Spalletti et al., 2014) that mimics a robot for upper limb rehabilitation in humans, the “Arm-Guide” (Reinkensmeyer et al., 2001). In Spalletti et al., 2017, robotic rehabilitation paradigm *per se* was not able to generalize recovery to untrained motor tasks (i.e. Schallert Cylinder test). They also showed that BoNT/E silencing, active in the motor cortex for about 10 days after injection, produced a small and transient improvement in general forelimb motor tests (Gridwalk and Schallert Cylinder test). Nevertheless, the guide of an appropriate motor rehabilitation regime was necessary to achieve a complete functional recovery. Here we show that incomplete motor recovery produced by the single treatments is associated with non-physiological synaptic turnover and reduced spine density. The synergy between BoNT/E treatment and robotic training on other hand was able to restore the stability and density of spines of pre-stroke conditions. Finally, our results show that the generalized recovery of motor function exclusively induced by the combined therapy was associated with increased vascular proliferation in the peri-infarct area.

A commonly agreed hypothesis is that the genesis of an enriched vascular milieu around the infarct area might promote synaptogenesis and dendritic remodeling, in addition to axonal sprouting (Brown et al., 2010; Carmichael, 2006; Kelley and Steward, 1997). In parallel, a recent review by Wahl and colleagues suggested that rehabilitative training might shape the spared and the new circuits by stabilizing the active contacts (Wahl et al., 2014). In support of these hypotheses, we show that augmented vascular density induced by the combined rehabilitative treatment is associated with increased density of synaptic contacts and stabilization of synaptic turnover. Although there are no other *in vivo* studies examining rehabilitation-induced spine plasticity after stroke to make a direct comparison with, our results are in agreement with recent postmortem histological studies where rehabilitation was shown to determine significant increases in spine density of distal apical dendrites in corticospinal neurons (Wang et al., 2016).

In this study, we have shown for the first time that rehabilitation promotes the recovery of structural features of healthy neuronal networks. Only the combined therapy could rescue the random orientation of pyramidal neurons while supporting the stabilization of synaptic contacts. Interestingly, in the same REHAB mice where stroke profoundly altered the spatial distribution of blood vessels around the core (Figure 3D), the randomness of dendrite orientation resembled healthy (CTRL) mice (Figure 2B). Further analysis on the acute phase after stroke will allow understanding if random neuronal orientation is maintained or later recovered by rehabilitation.

We hypothesized that the neural plasticity we observed in rehabilitated mice could be supported by revascularization. This assumption is based on the relationship existing between the proangiogenic state and neurological improvement in patients with stroke (Arkuszewski et al., 2009; Hermann and Chopp, 2012; Xiong et al., 2010). The reason behind it could be that the region of active neovascularization acts as a niche for enhanced neuronal remodeling and tissue repair (Prakash and Carmichael, 2015; Shen et al., 2008). Here, we found that vascular density in the peri-infarct area is comparable to healthy animals in stroke non-treated mice (STROKE) and single treatment groups (TOXIN, ROBOT), while exceed physiological values in mice under double treatment (REHAB). Our results suggest that the transient proangiogenic state induced in response to an ischemic insult is enhanced by combined rehabilitation. This hypothesis is supported by our findings on increased endothelial cell proliferation in REHAB mice. In conjunction with the more efficient neuronal activation and stabilized synaptic turnover we measured in the same areas, our study thus supports the hypothesis that enduring recovery from stroke might result from the association of angiogenesis with neuronal plasticity (Ergul et al., 2012).

We then investigated the changes induced by the combined rehabilitation paradigm on cortical activity. It has been reported that motor-targeted focal stroke induces abnormally scattered cortical maps that persist for months after stroke (see (Harrison et al., 2013)). In our study, the diffuse structure of motor representation was observed in non-treated animals (STROKE group). The combined rehabilitation paradigm progressively re-established a cluster of neuronal activity in peri-infarct areas where location, timing and amplitude parameters highly resembled those of healthy control animals. Several studies recently addressed the contamination of the blood volume changes on fluorescence signal of calcium indicators like GCaMP (see, for instance, (Ma et al., 2016a; Ma et al., 2016b; Makino et al., 2017; Murphy et al., 2018; Wright et al., 2017)). The recent paper by Ma et al. (Ma et al., 2016b) showed that the hemodynamic contribution is slower than the GCaMP6f signal, with an average temporal delay of 0.86 ± 0.05 s (representing the phase shift between neural activity and total hemoglobin concentration). More recently, by simultaneously recording (wide-field) calcium dynamics and hemodynamics, Wright et al. (Wright et al., 2017) showed that sensory evoked responses corresponding to oxygenated and deoxygenated hemoglobin have (i) a very slow onset after stimulus presentation, (ii) delayed peaks and (iii) low peak magnitudes (see also (Murphy et al., 2016)) compared to GCaMP6 fluorescence. These studies suggest that the hemodynamic fluctuations, much slower than calcium-associated neuronal activity, affect the late phase of calcium transients more than the rise. Last, by performing wide-field imaging on awake Thy1-GFP mice, Makino an colleagues (Makino et al., 2017) confirmed that the majority of movement-evoked fluorescence changes recorded in Thy1-GCaMP6s mice are due to calcium rather than hemodynamic signals or other artifacts. In our study, we focused on the analysis of the rise phase of the calcium transient (i.e. slope and amplitude), where the hemodynamic contamination should be very low. Nevertheless, we cannot exclude that hemodynamic fluctuations could give a small contribution to the fluorescence signal we recorded on GCaMP6f mice.

By performing optogenetic stimulation on the same group of stroked-afflicted, rehabilitated mice, we demonstrated that the progressive spatial and temporal refocusing of motor control in the ipsilesional cortex is associated with increased interhemispheric connectivity. The functional coupling between homotopic motor cortices was restored 4 weeks after injury in STROKE mice, suggesting that some form of spontaneous recovery compensated for the transcallosal projections lost after stroke. The interhemispheric connectivity is further enhanced after the rehabilitative therapy, suggesting a possible correlation between the recovery of activation patterns in the perilesional area and the reinforcement of connectivity with pyramidal neurons projecting from healthy M1. While there are no studies examining, on the same animals, cortical functionality and transcallosal connectivity to draw direct comparisons with, our results are consistent with the hypothesis that rehabilitative training of the paretic upper limb can, on one side, increase the functional activation of motor regions of the ipsilesional cortex (Hodics et al., 2006; Hubbard et al., 2015) and, on the other, decrease interhemispheric inhibition from the contralesional motor cortex (Harris-Love et al., 2011). We speculate that refocusing of the motor representation could be supported by dendritic rewiring in the peri-lesional area (as shown by our *in vivo* imaging result on structural plasticity), as well as by the activity of contralesional pyramidal cells projecting to the peri-infarct area. Further analysis on targeted neuronal populations is needed to clarify how rehabilitation-induced changes in transcallosal connectivity are specifically linked to alterations in the excitatory/inhibitory balance in the ipsilesional spared tissue.

The mouse model of stroke used in this study provides several advantages over more traditional models. Recently, Lim and colleagues used voltage-sensitive dyes combined with optogenetics to demonstrate a spontaneous partial recovery of cortical functional connectivity 8 weeks after stroke (Lim et al., 2014). The combination of transgenic GCaMP6 mice with AAV-induced expression of ChR2 in the homotopic M1, has the advantage of enabling on one side, the reliable control and on the other, the stable monitoring of neuronal activity over weeks and months. We anticipate that the all-optical approach we used will be further extended to the longitudinal exploration of the modified transcallosal axonal projections from the injured to the heathy hemisphere in our rehabilitation paradigm.

Our multi-scale investigation brought to light complementary aspects of the structural and functional plasticity induced by rehabilitation that may lead to the development of more efficient therapies and improve post-stroke recovery in patient populations.

## Experimental procedures

### Animal and surgery

Animal protocols were approved by the Italian Minister of Health. GFP transgenic mice were implanted with a cranial window in the right hemisphere from the motor cortex up to the visual area. A thinned skull preparation was used on GCaMP6f mice. Each group contained comparable numbers of male and female mice, and the age of mice was consistent between the groups (4-12 months). The surgery was followed by the first imaging session under the two-photon microscope.

### Photothrombotic Stroke

Mice were injected with a Rose Bengal solution (0.2 ml, 10 mg/ml solution in Phosphate Buffer Saline (PBS)). Five minutes after intraperitoneal injection a white light from an LED lamp was focused with a 20X objective and used to illuminate the M1 for 15 min to induce unilateral stroke in the right hemisphere. During the same procedure, on REHAB mice we injected 500 nl of BoNT/E (80 nM) divided in 2 separate injections of 250 nl at (i) +0.5 anteroposterior, +1.75 mediolateral and (ii) +0.4 anteroposterior, +1.75 mediolateral at 700 μm cortical depth. For optogenetics experiments, we delivered 1 μl of AAV9-CaMKII-ChR2-mCherry (2.48*1013 GC/mL) 700-900 μm deep inside the cortex (Conti et al., 2019).

### Robotic Platform

The animals were trained by means of the M-Platform (Spalletti et al., 2014), which is a robotic system that allows mice to perform a retraction movement of their left forelimb. The motor rehabilitation consists in a pulling task: first the animal forelimb is passively extended by the linear actuator of the platform and then the animal has to pull back the forelimb up to the resting position. The motor training is composed by 15 movements and after each movement the animal receives a liquid reward.

### Labelling of brain vasculature

After perfusion with PFA 4% followed by a perfusion with 10 ml of fluorescent gel, the vasculature was stained using the protocol described by Tsai et al.(Tsai et al., 2009), except that we replaced fluorescein (FITC)-conjugated albumin with 0.05% (w/v) tetramethylrhodamine (TRITC)-conjugated albumin in order to avoid spectral overlap between GFP and FITC (Di Giovanna et al., 2018).

### Statistical Analysis

Data were analyzed using Origin Pro. The specific tests used are stated alongside all probability values reported.

## Acknowledgement

We thank Alessio Masi and Marie Caroline Muellenbroich for very useful discussion about the manuscript; Giuseppe De Vito for assistance on statistics analysis. We thank Dr. Zanier from the Mario Negri Institute (Milan) for providing reagents for immunohistochemical analysis. We thank the mechanics and electronics workshops at LENS. This project has received funding from the H2020 EXCELLENT SCIENCE -European Research Council (ERC) under grant agreement ID n. 692943 BrainBIT”. In addition, it was supported by the European Union’s Horizon 2020 research and innovation program under grant agreements No. 720270 (Human Brain Project) and 654148 (Laserlab-Europe). Part of this work was performed within the framework of the Proof of Concept Studies for the ESFRI research infrastructure project Euro-BioImaging at the PCS facility LENS.

## Author Contributions

A.L.A.M. and M.C. conceived the study. A.L.A.M., E.C., A.P.D., C.S and C.A. performed experiments. S.L., A.L.A.M., E.C., A.S. and A.P.D. processed data. A.P., S.L. and L.S. developed the integrated microscope and M-platform system. F.S.P., S.M. and M.C. obtained funding support. A.L.A.M. and E.C. wrote the paper. All authors approved the paper.

**Declaration of Interests**

The authors declare no competing interests.

## References

Adkins-Muir, D.L., and Jones, T.A. (2003). Cortical electrical stimulation combined with rehabilitative training: enhanced functional recovery and dendritic plasticity following focal cortical ischemia in rats. Neurological research 25, 780–788.

Ankarcrona, M., Dypbukt, J.M., Bonfoco, E., Zhivotovsky, B., Orrenius, S., Lipton, S.A., and Nicotera, P. (1995). Glutamate-induced neuronal death: a succession of necrosis or apoptosis depending on mitochondrial function. Neuron 15, 961–973.

Arkuszewski, M., Świat, M., and Opala, G. (2009). Perfusion computed tomography in prediction of functional outcome in patients with acute ischaemic stroke. Nuclear Medicine Review 12, 89–94.

Ayling, O.G., Harrison, T.C., Boyd, J.D., Goroshkov, A., and Murphy, T.H. (2009). Automated light-based mapping of motor cortex by photoactivation of channelrhodopsin-2 transgenic mice. Nature methods 6, 219–224.

Bauer, A.Q., Kraft, A.W., Wright, P.W., Snyder, A.Z., Lee, J.M., and Culver, J.P. (2014). Optical imaging of disrupted functional connectivity following ischemic stroke in mice. NeuroImage 99, 388–401.

Brown, C.E., Aminoltejari, K., Erb, H., Winship, I.R., and Murphy, T.H. (2009). In vivo voltage-sensitive dye imaging in adult mice reveals that somatosensory maps lost to stroke are replaced over weeks by new structural and functional circuits with prolonged modes of activation within both the peri-infarct zone and distant sites. The Journal of neuroscience : the official journal of the Society for Neuroscience 29, 1719–1734.

Brown, C.E., Boyd, J.D., and Murphy, T.H. (2010). Longitudinal in vivo imaging reveals balanced and branch-specific remodeling of mature cortical pyramidal dendritic arbors after stroke. Journal of cerebral blood flow and metabolism : official journal of the International Society of Cerebral Blood Flow and Metabolism 30, 783–791.

Brown, C.E., Li, P., Boyd, J.D., Delaney, K.R., and Murphy, T.H. (2007a). Extensive turnover of dendritic spines and vascular remodeling in cortical tissues recovering from stroke. J Neurosci 27, 4101–4109.

Brown, C.E., Li, P., Boyd, J.D., Delaney, K.R., and Murphy, T.H. (2007b). Extensive Turnover of Dendritic Spines and Vascular Remodeling in Cortical Tissues Recovering from Stroke. The Journal of Neuroscience 27, 4101–4109.

Bütefisch, C.M. (2006). Neurobiological bases of rehabilitation. Neurological Sciences 27, s18–s23.

Caleo, M., Restani, L., Gianfranceschi, L., Costantin, L., Rossi, C., Rossetto, O., Montecucco, C., and Maffei, L. (2007). Transient synaptic silencing of developing striate cortex has persistent effects on visual function and plasticity. The Journal of neuroscience : the official journal of the Society for Neuroscience 27, 4530–4540.

Carmichael, S.T. (2006). Cellular and molecular mechanisms of neural repair after stroke: making waves. Annals of neurology 59, 735–742.

Carmichael, S.T., Kathirvelu, B., Schweppe, C.A., and Nie, E.H. (2017). Molecular, cellular and functional events in axonal sprouting after stroke. Experimental neurology 287, 384–394.

Carmichael, S.T., Wei, L., Rovainen, C.M., and Woolsey, T.A. (2001). New patterns of intracortical projections after focal cortical stroke. Neurobiology of disease 8, 910–922.

Carter, A.R., Astafiev, S.V., Lang, C.E., Connor, L.T., Rengachary, J., Strube, M.J., Pope, D.L., Shulman, G.L., and Corbetta, M. (2010). Resting interhemispheric functional magnetic resonance imaging connectivity predicts performance after stroke. Annals of neurology 67, 365–375.

Chen, T.-W., Wardill, T.J., Sun, Y., Pulver, S.R., Renninger, S.L., Baohan, A., Schreiter, E.R., Kerr, R.A., Orger, M.B., Jayaraman, V., et al. (2013). Ultrasensitive fluorescent proteins for imaging neuronal activity. Nature 499, 295–300.

Clarkson, A.N., Huang, B.S., Macisaac, S.E., Mody, I., and Carmichael, S.T. (2010). Reducing excessive GABA-mediated tonic inhibition promotes functional recovery after stroke. Nature 468, 305–309.

Conti, E., Allegra Mascaro, A.L., and Pavone, F.S. (2019). Large Scale Double-Path Illumination System with Split Field of View for the All-Optical Study of Inter-and Intra-Hemispheric Functional Connectivity on Mice. Methods and Protocols 2, 11.

Crocini, C., Ferrantini, C., Coppini, R., Scardigli, M., Yan, P., Loew, L.M., Smith, G., Cerbai, E., Poggesi, C., Pavone, F.S., et al. (2016). Optogenetics design of mechanistically-based stimulation patterns for cardiac defibrillation. Scientific reports 6, 35628.

Dancause, N., Barbay, S., Frost, S.B., Plautz, E.J., Chen, D., Zoubina, E.V., Stowe, A.M., and Nudo, R.J. (2005). Extensive cortical rewiring after brain injury. The Journal of neuroscience : the official journal of the Society for Neuroscience 25, 10167–10179.

Dancause, N., and Nudo, R.J. (2011). Shaping plasticity to enhance recovery after injury. Progress in brain research 192, 273–295.

Di Giovanna, A.P., Tibo, A., Silvestri, L., Müllenbroich, M.C., Costantini, I., Allegra Mascaro, A.L., Sacconi, L., Frasconi, P., and Pavone, F.S. (2018). Whole-Brain Vasculature Reconstruction at the Single Capillary Level. Scientific Reports 8, 12573.

Ergul, A., Alhusban, A., and Fagan, S.C. (2012). Angiogenesis: A Harmonized Target for Recovery after Stroke. Stroke 43, 2270–2274.

Fang, P.C., Barbay, S., Plautz, E.J., Hoover, E., Strittmatter, S.M., and Nudo, R.J. (2010). Combination of NEP 1-40 treatment and motor training enhances behavioral recovery after a focal cortical infarct in rats. Strok 41, 544–549.

Harris-Love, M.L., Morton, S.M., Perez, M.A., and Cohen, L.G. (2011). Mechanisms of short-term training-induced reaching improvement in severely hemiparetic stroke patients: a TMS study. Neurorehabilitation and neural repair 25, 398–411.

Harrison, T.C., Silasi, G., Boyd, J.D., and Murphy, T.H. (2013). Displacement of sensory maps and disorganization of motor cortex after targeted stroke in mice. Stroke 44, 2300–2306.

Hermann, D.M., and Chopp, M. (2012). Promoting brain remodelling and plasticity for stroke recovery: therapeutic promise and potential pitfalls of clinical translation. The Lancet Neurology 11, 369–380.

Hesse, S., Werner, C., Schonhardt, E.M., Bardeleben, A., Jenrich, W., and Kirker, S.G. (2007). Combined transcranial direct current stimulation and robot-assisted arm training in subacute stroke patients: a pilot study. Restorative neurology and neuroscience 25, 9–15.

Hodics, T., Cohen, L.G., and Cramer, S.C. (2006). Functional imaging of intervention effects in stroke motor rehabilitation. Archives of physical medicine and rehabilitation 87, S36–42.

Hsu, J.E., and Jones, T.A. (2006). Contralesional neural plasticity and functional changes in the less-affected forelimb after large and small cortical infarcts in rats. Experimental neurology 201, 479–494.

Hubbard, I.J., Carey, L.M., Budd, T.W., Levi, C., McElduff, P., Hudson, S., Bateman, G., and Parsons, M.W. (2015). A Randomized Controlled Trial of the Effect of Early Upper-Limb Training on Stroke Recovery and Brain Activation. Neurorehabilitation and neural repair 29, 703–713.

Johnston, D.G., Denizet, M., Mostany, R., and Portera-Cailliau, C. (2013). Chronic in vivo imaging shows no evidence of dendritic plasticity or functional remapping in the contralesional cortex after stroke. Cerebral cortex 23, 751–762.

Jones, T.A., and Adkins, D.L. (2015). Motor System Reorganization After Stroke: Stimulating and Training Toward Perfection. Physiology 30, 358–370.

Kelley, M.S., and Steward, O. (1997). Injury-induced physiological events that may modulate gene expression in neurons and glia. Reviews in the neurosciences 8, 147–177.

Krakauer, J.W., Carmichael, S.T., Corbett, D., and Wittenberg, G.F. (2012). Getting neurorehabilitation right: what can be learned from animal models? Neurorehabilitation and neural repair 26, 923–931.

Lee, J.K., Kim, J.E., Sivula, M., and Strittmatter, S.M. (2004). Nogo receptor antagonism promotes stroke recovery by enhancing axonal plasticity. The Journal of neuroscience : the official journal of the Society for Neuroscience 24, 6209–6217.

Lim, D.H., LeDue, J.M., Mohajerani, M.H., and Murphy, T.H. (2014). Optogenetic mapping after stroke reveals network-wide scaling of functional connections and heterogeneous recovery of the peri-infarct. The Journal of neuroscience : the official journal of the Society for Neuroscience 34, 16455–16466.

Ma, Y., Shaik, M.A., Kim, S.H., Kozberg, M.G., Thibodeaux, D.N., Zhao, H.T., Yu, H., and Hillman, E.M. (2016a). Wide-field optical mapping of neural activity and brain haemodynamics: considerations and novel approaches. Philosophical Transactions of the Royal Society B: Biological Sciences 371, 20150360.

Ma, Y., Shaik, M.A., Kozberg, M.G., Kim, S.H., Portes, J.P., Timerman, D., and Hillman, E.M. (2016b). Resting-state hemodynamics are spatiotemporally coupled to synchronized and symmetric neural activity in excitatory neurons. Proceedings of the National Academy of Sciences of the United States of America 113, E8463–E8471.

Makino, H., Ren, C., Liu, H., Kim, A.N., Kondapaneni, N., Liu, X., Kuzum, D., and Komiyama, T. (2017). Transformation of Cortex-wide Emergent Properties during Motor Learning. Neuron 94, 880–890 e888.

Mostany, R., Chowdhury, T.G., Johnston, D.G., Portonovo, S.A., Carmichael, S.T., and Portera-Cailliau, C. (2010). Local hemodynamics dictate long-term dendritic plasticity in peri-infarct cortex. The Journal of neuroscience : the official journal of the Society for Neuroscience 30, 14116–14126.

Mozaffarian, D., Benjamin, E.J., Go, A.S., Arnett, D.K., Blaha, M.J., Cushman, M., de Ferranti, S., Despres, J.P., Fullerton, H.J., Howard, V.J., et al. (2015). Heart disease and stroke statistics--2015 update: a report from the American Heart Association. Circulation 131, e29–322.

Murphy, M.C., Chan, K.C., Kim, S.G., and Vazquez, A.L. (2018). Macroscale variation in resting-state neuronal activity and connectivity assessed by simultaneous calcium imaging, hemodynamic imaging and electrophysiology. NeuroImage 169, 352–362.

Murphy, T.H., Boyd, J.D., Bolaños, F., Vanni, M.P., Silasi, G., Haupt, D., and LeDue, J.M. (2016). High-throughput automated home-cage mesoscopic functional imaging of mouse cortex. Nature Communications 7, 11611.

Plautz, E.J., Barbay, S., Frost, S.B., Friel, K.M., Dancause, N., Zoubina, E.V., Stowe, A.M., Quaney, B.M., and Nudo, R.J. (2003). Post-infarct cortical plasticity and behavioral recovery using concurrent cortical stimulation and rehabilitative training: a feasibility study in primates. Neurological research 25, 801–810.

Prakash, R., and Carmichael, S.T. (2015). Blood-brain barrier breakdown and neovascularization processes after stroke and traumatic brain injury. Current opinion in neurology 28, 556–564.

Reinkensmeyer, D.J., Takahashi, C.D., Timoszyk, W.K., Reinkensmeyer, A.N., and Kahn, L.E. (2001). Design of robot assistance for arm movement therapy following stroke. Advanced Robotics 14, 625–637.

Sakadzic, S., Lee, J., Boas, D.A., and Ayata, C. (2015). High-resolution in vivo optical imaging of stroke injury and repair. Brain research 1623, 174–192.

Shen, Q., Wang, Y., Kokovay, E., Lin, G., Chuang, S.M., Goderie, S.K., Roysam, B., and Temple, S. (2008). Adult SVZ stem cells lie in a vascular niche: a quantitative analysis of niche cell-cell interactions. Cell stem cell 3, 289–300.

Sigler, A., and Murphy, T.H. (2010). In vivo 2-photon imaging of fine structure in the rodent brain: before, during, and after stroke. Stroke 41, S117–123.

Spalletti, C., Alia, C., Lai, S., Panarese, A., Conti, S., Micera, S., and Caleo, M. (2017). Combining robotic training and inactivation of the healthy hemisphere restores pre-stroke motor patterns in mice. Elife 6, e28662.

Spalletti, C., Lai, S., Mainardi, M., Panarese, A., Ghionzoli, A., Alia, C., Gianfranceschi, L., Chisari, C., Micera, S., and Caleo, M. (2014). A robotic system for quantitative assessment and poststroke training of forelimb retraction in mice. Neurorehabilitation and neural repair 28, 188–196.

Tsai, P.S., Kaufhold, J.P., Blinder, P., Friedman, B., Drew, P.J., Karten, H.J., Lyden, P.D., and Kleinfeld,D. (2009). Correlations of neuronal and microvascular densities in murine cortex revealed by direct counting and colocalization of nuclei and vessels. The Journal of neuroscience : the official journal of the Society for Neuroscience 29, 14553–14570.

Ueno, Y., Chopp, M., Zhang, L., Buller, B., Liu, Z., Lehman, N.L., Liu, X.S., Zhang, Y., Roberts, C., and Zhang, Z.G. (2012). Axonal outgrowth and dendritic plasticity in the cortical peri-infarct area after experimental stroke. Stroke 43, 2221–2228.

van Meer, M.P., van der Marel, K., Otte, W.M., Berkelbach van der Sprenkel, J.W., and Dijkhuizen, R.M. (2010a). Correspondence between altered functional and structural connectivity in the contralesional sensorimotor cortex after unilateral stroke in rats: a combined resting-state functional MRI and manganese-enhanced MRI study. Journal of cerebral blood flow and metabolism : official journal of the International Society of Cerebral Blood Flow and Metabolism 30, 1707–1711.

van Meer, M.P., van der Marel, K., Wang, K., Otte, W.M., El Bouazati, S., Roeling, T.A., Viergever, M.A., Berkelbach van der Sprenkel, J.W., and Dijkhuizen, R.M. (2010b). Recovery of sensorimotor function after experimental stroke correlates with restoration of resting-state interhemispheric functional connectivity. The Journal of neuroscience : the official journal of the Society for Neuroscience 30, 3964–3972.

Wahl, A.S., Omlor, W., Rubio, J.C., Chen, J.L., Zheng, H., Schroter, A., Gullo, M., Weinmann, O., Kobayashi, K., Helmchen, F., et al. (2014). Neuronal repair. Asynchronous therapy restores motor control by rewiring of the rat corticospinal tract after stroke. Science 344, 1250–1255.

Wang, L., Conner, J.M., Nagahara, A.H., and Tuszynski, M.H. (2016). Rehabilitation drives enhancement of neuronal structure in functionally relevant neuronal subsets. Proceedings of the National Academy of Sciences of the United States of America 113, 2750–2755.

Wright, P.W., Brier, L.M., Bauer, A.Q., Baxter, G.A., Kraft, A.W., Reisman, M.D., Bice, A.R., Snyder, A.Z., Lee, J.M., and Culver, J.P. (2017). Functional connectivity structure of cortical calcium dynamics in anesthetized and awake mice. PloS one 12, e0185759.

Xiong, Y., Mahmood, A., and Chopp, M. (2010). Angiogenesis, neurogenesis and brain recovery of function following injury. Curr Opin Invest Dr 11, 298–308.

