## Supplementary figures and images for "Rehabilitation promotes the recovery of structural and functional features of healthy neuronal networks after stroke"

### Figure S1

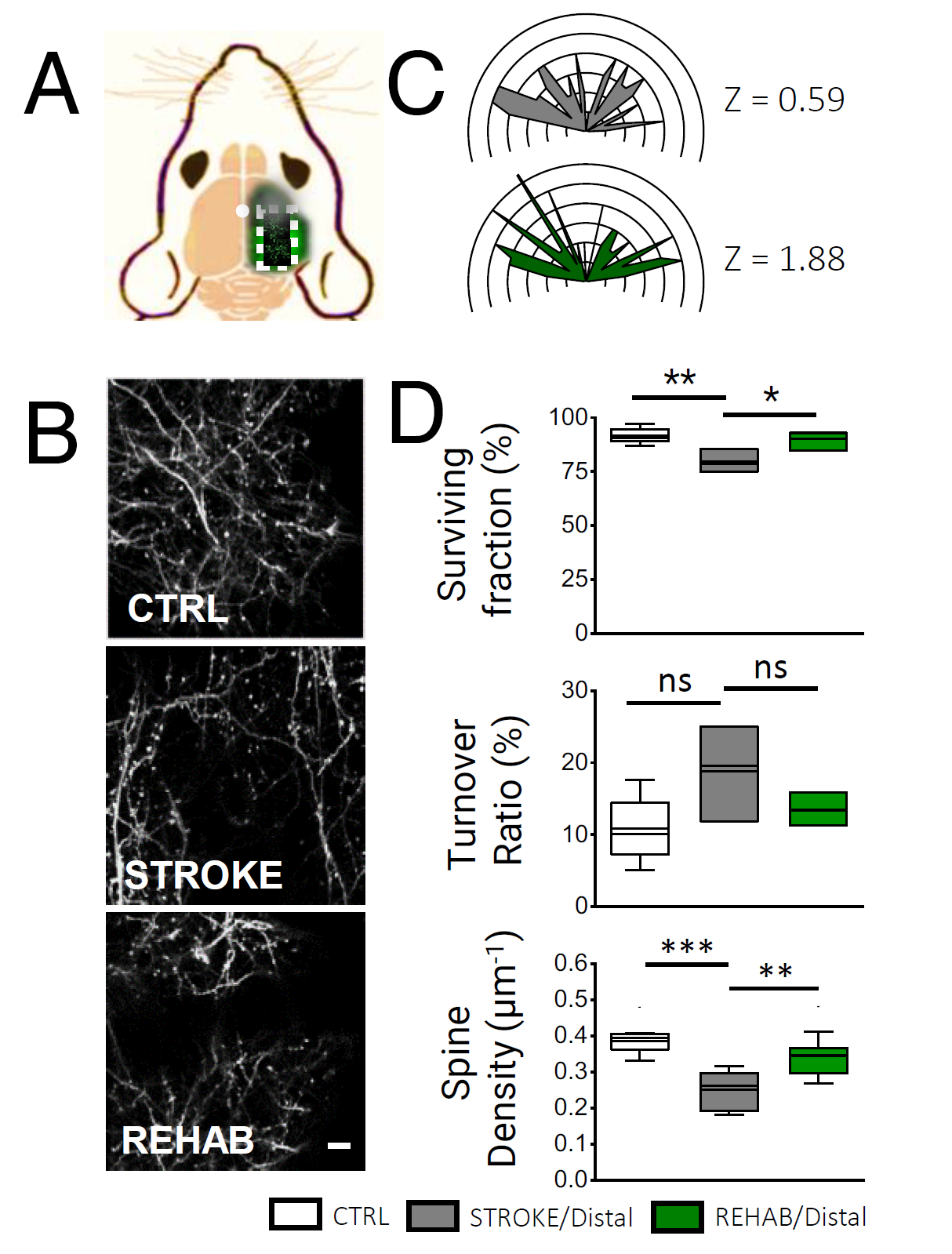

### Figure S2

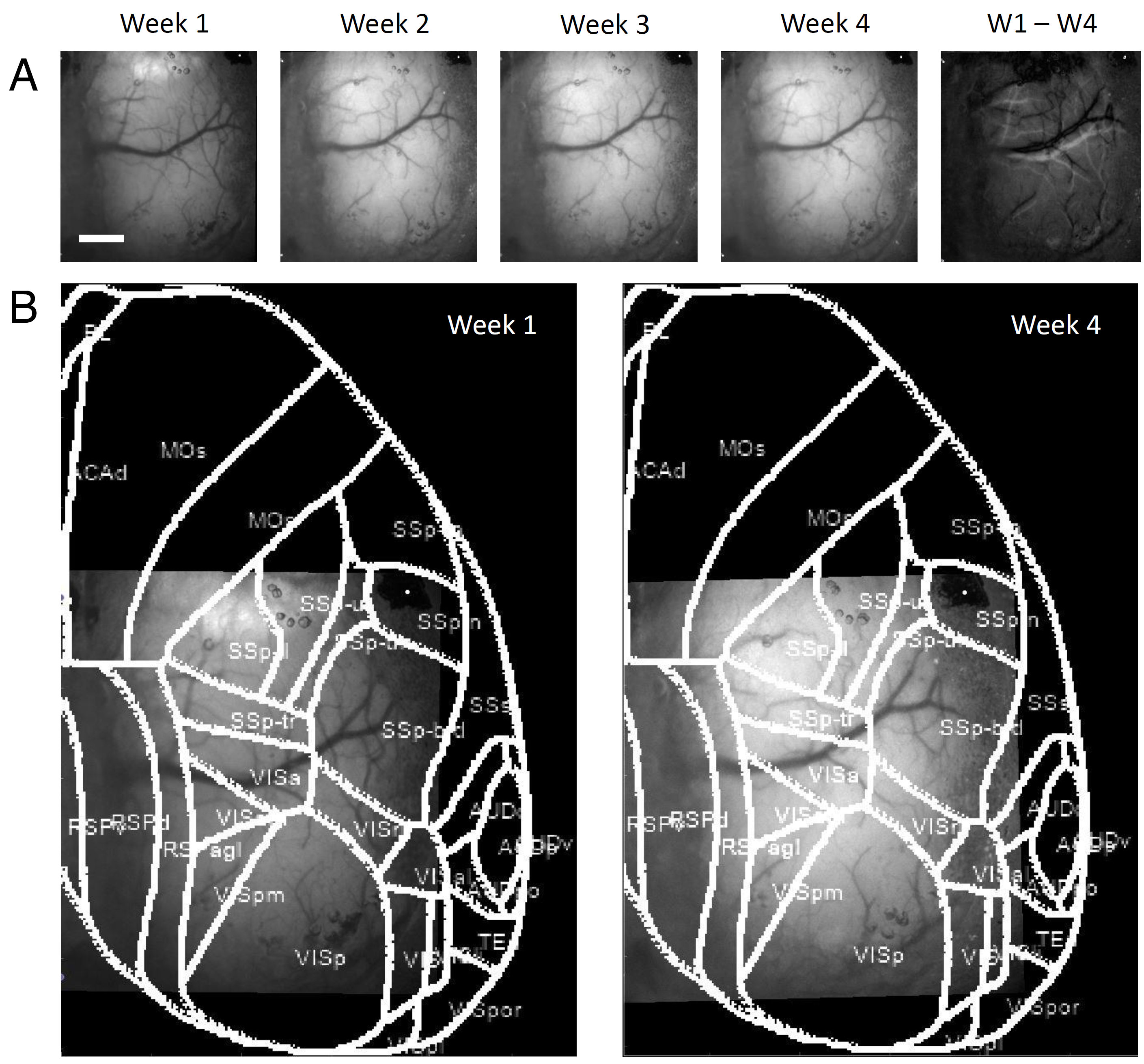

### Figure S3

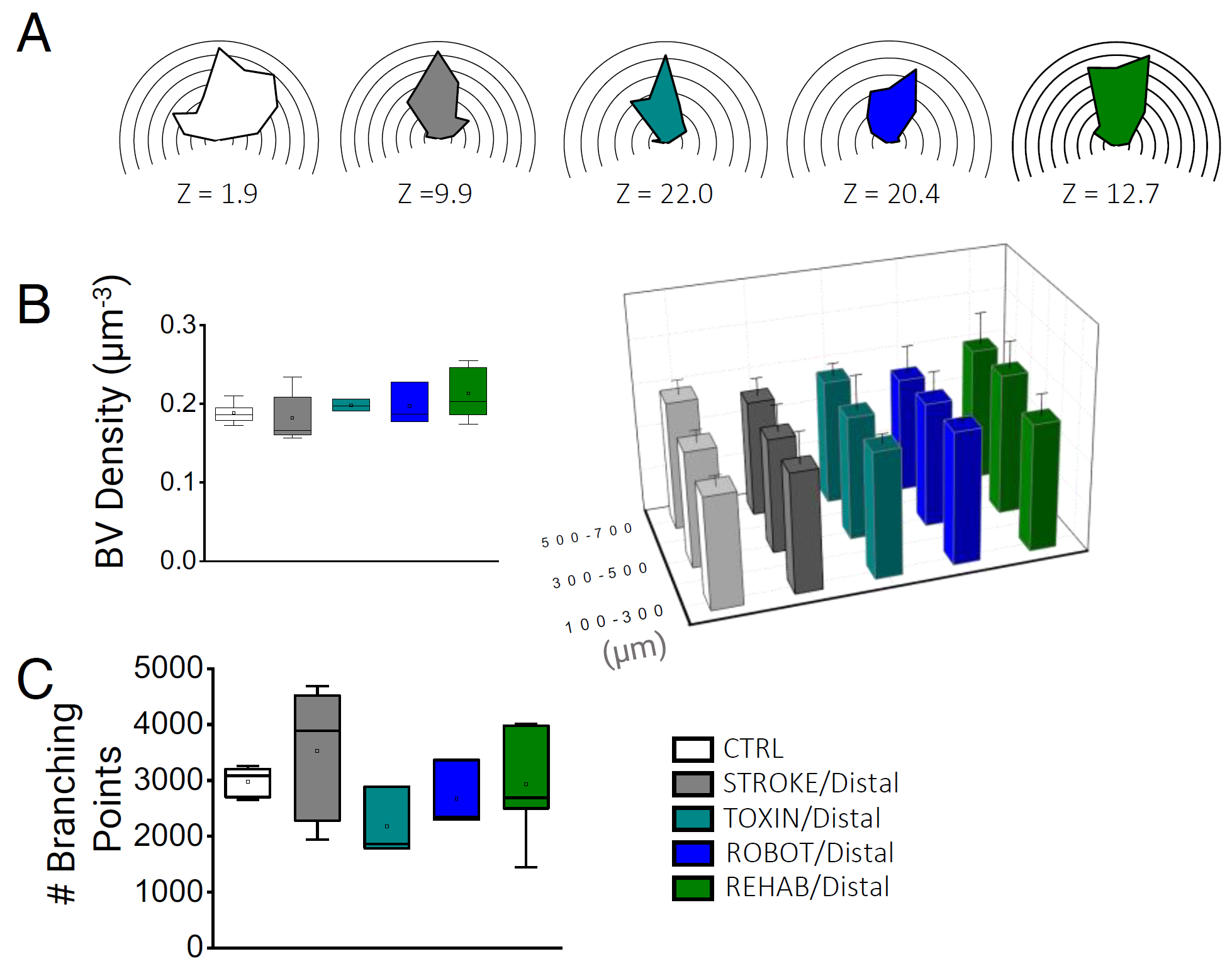

### Figure S4

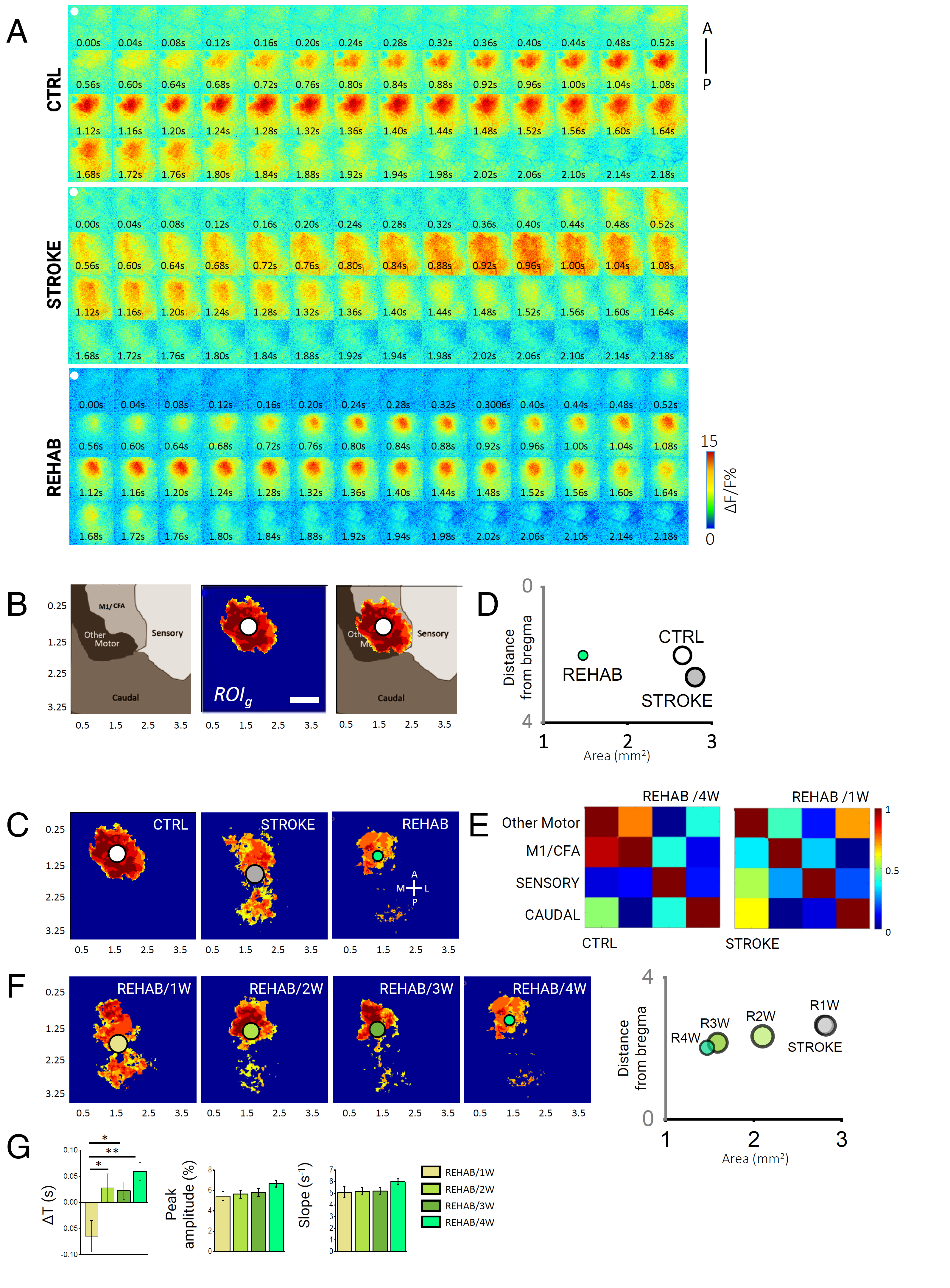

### Figure S5

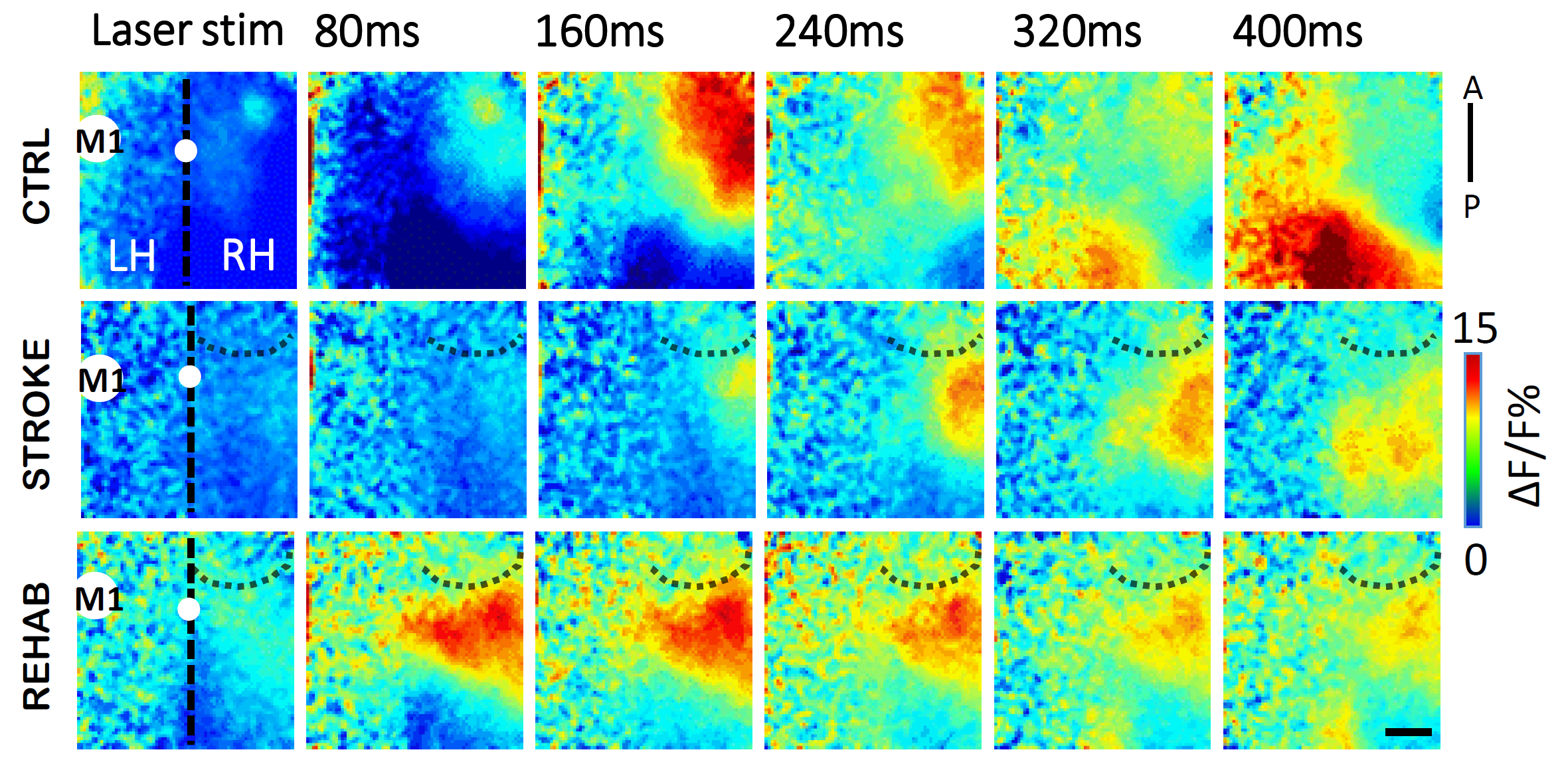
