## Supplementary material for "Rehabilitation promotes the recovery of structural and functional features of healthy neuronal networks after stroke"

### Supplemental Figures

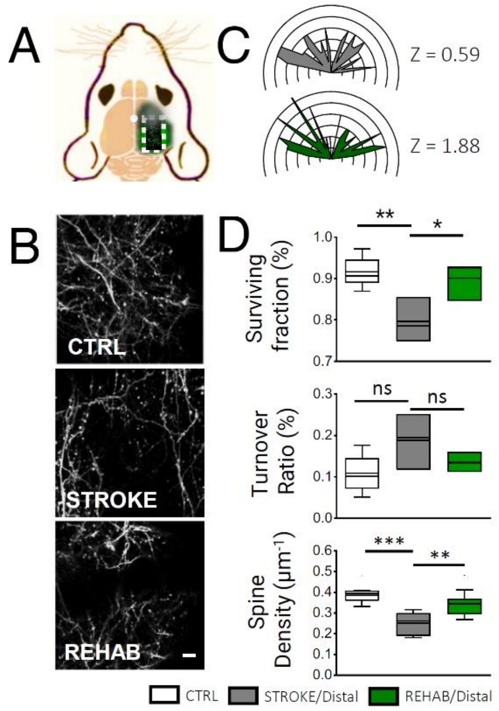


Figure S1. In vivo imaging of dendritic and spine plasticity in regions proximal and distal from stroke core. Related to Figure 2. (A) A schematic representation of the field of view (i.e. area within the white dotted square) of two-photon imaging of dendritic and spine plasticity in GFPM mice. The red spot indicates the site of the stroke lesion. (B) Representative examples of maximum intensity projection of two-photons stacks of dendritic branch orientation in the peri-infarct area in the CTRL, STROKE and REHAB groups. Scale bar, 10 µm. (C) Polar plots showing the angular distribution of dendrites in the distal area ($Nmice_{CTRL}$= 5, $Nmice_{STROKE}$ = 4, $Nmice_{REHAB}$= 3). Z scores, calculated by the Rayleigh Test for circular statistics, are reported for each experimental class on the right of the polar plot; ns, not significant). (D) Histograms showing the SF (*upper panel;* $Nmice_{CTRL}$= 6, $Nmice_{STROKE}$ = 4, $Nmice_{REHAB}$= 3; $SF_{CTRL}$= 0.92 ± 0.02%; $SF_{STROKE\ Distal}$= 0.80 ± 0.03%; $SF_{REHAB\ DIstal}$= 0.90 ± 0.03%; one way ANOVA with post hoc Bonferroni: ** $P$ =0.007, * $P$ =0.023), TOR (*lower panel;* $Nmice_{CTRL}$= 6, $Nmice_{STROKE}$ = 4, $Nmice_{REHAB}$= 3; $TOR_{CTRL}$= 0.11 ± 0.02%; $TOR_{STROKE\ Distal}$ = 0.19 ± 0.04%; $TOR_{REHAB}$= 0.13 ± 0.01% t-test: one way ANOVA with post hoc Bonferroni: not significant for all comparisons) and spine density (SD) (*right panel*; $Nmice_{CTRL}$= 3, $Nmice_{STROKE}$ = 3, $Nmice_{REHAB}$= 3; $SD_{CTRL}$= 0.39 ± 0.01 $\mu m^{-1}$; $SD_{STROKE}$= 0.25 ± 0.02 $\mu m^{-1}$; $SD_{REHAB}$= 0.35 ± 0.02 $\mu m^{-1}$; one-way ANOVA with post hoc Fisher test, $P_{CTRL/STROKE}$ = 0.0001, $P_{REHAB/STROKE}$ = 0.005) in the distal region (>1000 µm from the stroke core). Data are means ± SEM.

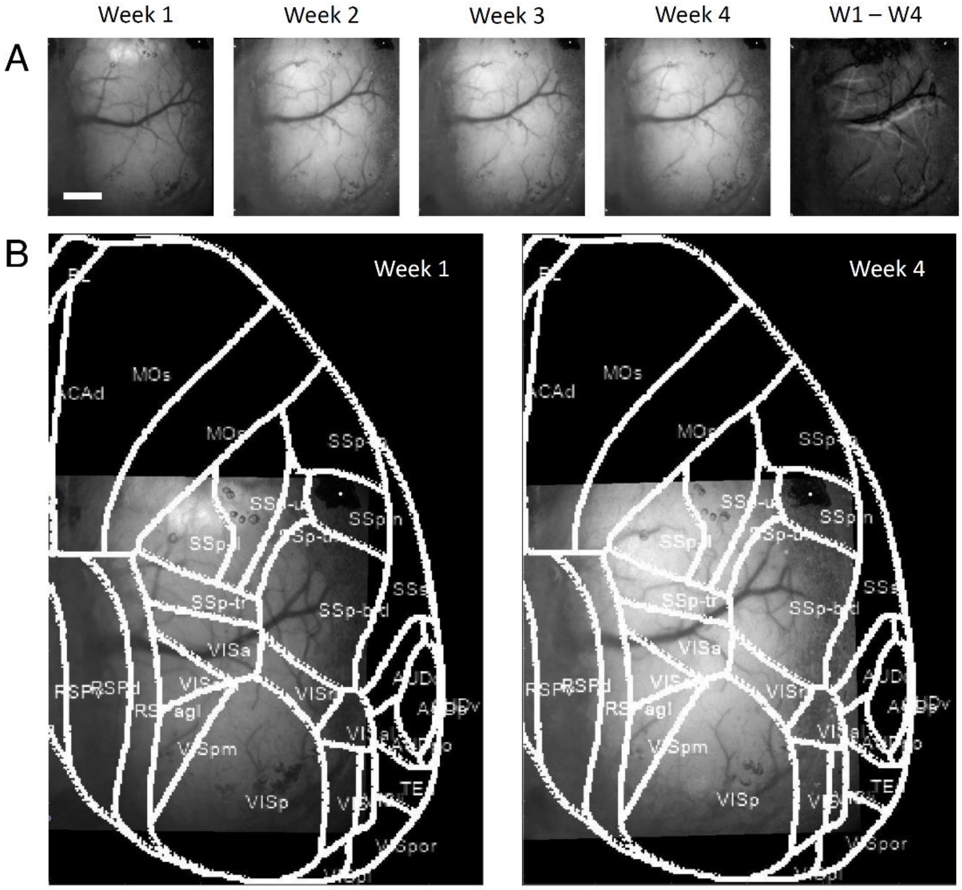


Figure S2. Shrinkage induced by cell necrosis in the stroke core progressively shifts cortical tissue towards the lesioned area. Related to Figure 3. (A) Example of longitudinal brightfield images acquired on every week after stroke showing progressive displacement of cortical vasculature. The last image on the right highlights the displacement of blood vessels between the 1st and 4th week after stroke. Scale bar, 1 mm (B) The same images taken on the 1st and 4th week after stroke are superimposed on the Allen Mouse Brain Atlas (https://mouse.brain-map.org/static/atlas) of the right hemisphere to better visualize the localization of our field of view and blood vessel displacement.

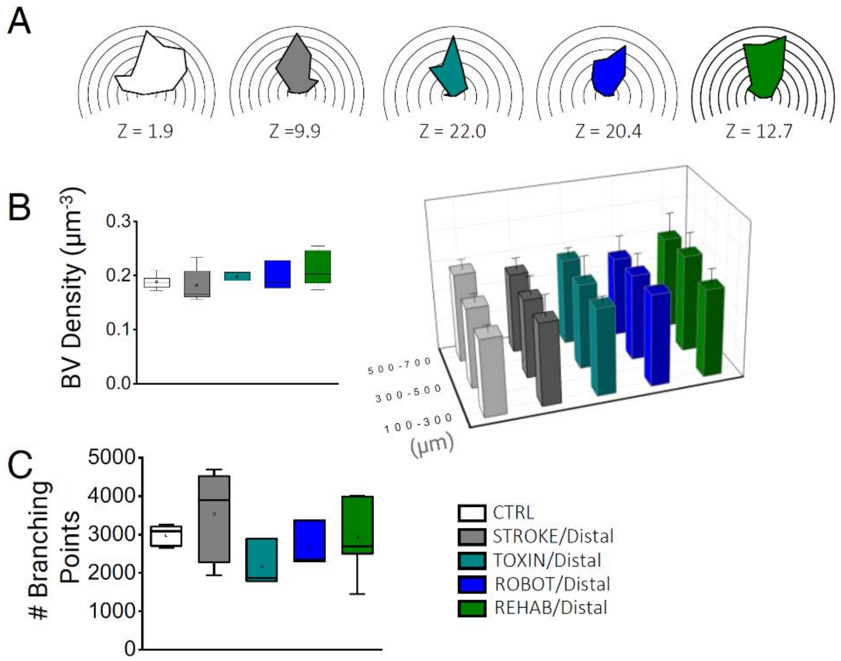


Figure S3. Distal analysis of blood vessels. Related to Figure 3. (A) Polar plots show the distribution of blood vessel orientation in regions distal to the core for each experimental group measured after the last motor training session ($Nmice_{CTRL}$ = 5; $Nmice_{STROKE}$ = 6; $Nmice_{TOXIN}$ = 3; $Nmice_{ROBOT}$ = 3; $Nmice_{REHAB}$ = 5). Z scores, calculated by the Rayleigh Test for circular statistics, are reported for each experimental class below the plot. (B) Blood vessel (BV) density analysis. *(Left)* The box and whiskers plot shows the BV density (average ± SEM) in regions distal from the stroke core in each experimental group (BV $Density_{CTRL}$ = 0.19 ± 0.01 $\mu m^{-3}$; BV $Density_{STROKE\ Distal}$ = 0.18 ± 0.03; BV $Density_{TOXIN\ Distal}$= 0.20 ± 0.01; BV $Density_{ROBOT\ Distal}$ = 0.20 ± 0.03; BV $Density_{REHAB\ Distal}$ = 0.21 ± 0.04; one-way ANOVA with post hoc Bonferroni test, n.s.). *(Right)* 3D graph comparing the average BV density grouped by cortical depth (100-300 µm from stroke core: BV $Density_{CTRL}$ = 0.18 ± 0.02; BV $Density_{STROKE\ Distal}$ = 0.19 ± 0.05; BV $Density_{TOXIN\ Distal}$ = 0.20 ± 0.01; BV $Density_{ROBOT\ Distal}$ = 0.21 ± 0.01; BV $Density_{REHAB\ Distal}$ = 0.20 ± 0.03; 300-500 µm from stroke core: BV $Density_{CTRL}$ = 0.18 ± 0.02; BV $Density_{STROKE\ Distal}$ = 0.18 ± 0.04; BV $Density_{TOXIN\ Distal}$ = 0.19 ± 0.05; BV $Density_{ROBOT\ Distal}$ = 0.20 ± 0.02; BV $Density_{REHAB\ Distal}$ = 0.22 ± 0.04; 500-700 µm from stroke core: BV $Density_{CTRL}$ = 0.20 ± 0.02; BV $Density_{STROKE\ Distal}$ = 0.18 ± 0.03; BV $Density_{TOXIN\ Distal}$ = 0.20 ± 0.02; BV $Density_{ROBOT\ Distal}$= 0.18 ± 0.04; BV $Density_{REHAB\ Distal}$ = 0.21 ± 0.05. (C) The box and whiskers plot shows the quantification of the number of branching points for each experimental group in distal regions from the stroke core (Number of Branching Points (BP): $BP_{CTRL}$ = 2979 ± 126; $BP_{STROKE}$ = 3533 ± 474; $BP_{TOXIN}$ = 2179 ± 355; $BP_{ROBOT}$ = 2670 ± 347; $BP_{REHAB}$ = 2927 ± 486).

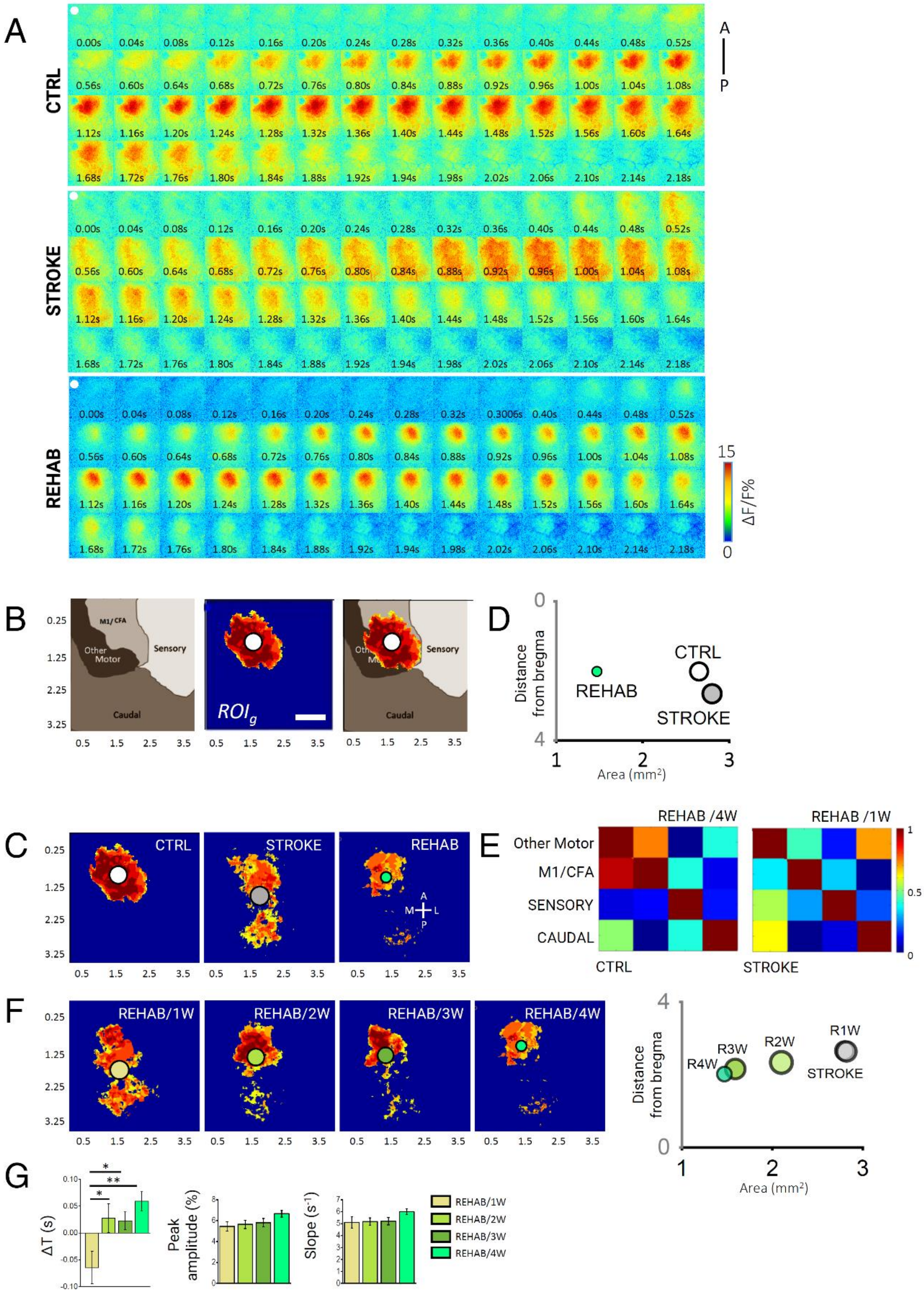


Figure S4. Wide-field imaging of cortical activation profiles over 4 weeks of rehabilitation. Related to Figure 4. (A) Complete image sequences of cortical activation as assessed by calcium imaging during pulling of the handle by the contralateral forelimb of CTRL (top), STROKE (middle) and REHAB (bottom) GCaMP6f mice in the M-Platform. A-P, anterior-posterior. The white dot indicates bregma. Scale bar = 1 mm. (B) The image on the left shows a functional map based on the intracortical microstimulation (ICMS) studies of Tennant (2011, Cerebral Cortex) and Alia (2016, Sci Reports) that was used as a reference map. The middle panel shows the average thresholded ROI ($ROI_g$) computed for CTRL animals during voluntary contralateral forelimb pulling in the M-Platform. The image on the right shows a merged image of the functional map and *ROIg*. CFA, caudal forelimb area. The caudal area includes visual and associative regions. Scale bar, 1 mm. (C) The panels show the average thresholded ROI ($ROI_g$) computed for each experimental group. The circles represent the centroids of the $ROI_g$ for CTRL (white), STROKE (light gray), and REHAB (green) groups. (D) The areas of the $ROI_g$ and their distance from bregma are presented in the scatter plot. (E) Examples of partial correlation matrices of cortical activation during voluntary pulling. The correlation analysis on REHAB mice was performed based on two evaluation time points: one week (REHAB/1W) and four weeks (REHAB/4W) after stroke. (F) On the left, average thresholded ROI ($ROI_g$) computed for each week of the REHAB group. The circles represent the centroids of the $ROI_g$ for REHAB/1W (light yellow), REHAB/2W (light green), REHAB/3W (olive green), REHAB/4W (bright green) groups. On the right, the areas of the $ROI_g$ and their distance from bregma are presented in the scatter plot for the 4 weeks of REHAB and the STROKE groups. (G) Longitudinal analysis of cortical activation profiles over the four rehabilitation weeks. Graphs in the left, middle and right panels are calculated as in Figure 4E, G, and H, respectively. Left panel: Delays in cortical activation in caudal regions following the forelimb retraction task are reported for the 4 weeks of rehabilitative training ($Nmice_{REHAB}$= 6; $\Delta T_{REHAB/1W}$ = -0.06 ± 0.03 s, $\Delta T_{REHAB/2W}$ = -0.03 ± 0.03 s, $\Delta T_{REHAB/3W}$ = 0.02 ± 0.02 s, $\Delta T_{REHAB/4W}$ = 0.06 ± 0.02 s; one-way ANOVA followed by the Tukey test: ** $P$ <0.01, *$P$ <0.05; REHAB/4W here corresponds to the REHAB group in all the other panels). Middle panel: the maximum of calcium imaging fluorescence peaks for the 4 weeks of rehabilitative training ($Nmice_{REHAB}$= 6; Peak $amplitude_{REHAB/1W}$= 5.4 ± 0.4%, Peak $amplitude_{REHAB/2W}$ = 5.6 ±0.4 %, Peak $amplitude_{REHAB/3W}$ = 5.8 ± 0.4%, Peak $amplitude_{REHAB/4W}$ = 6.6 ± 0.3 %; No significant changes were observed over the 4 weeks period). Right panel: The graph shows the slope (average ± SEM) of the calcium imaging fluorescence in the rising phase ($Nmice_{REHAB}$= 6; $Slope_{REHAB/1W}$= 5.1 ±0.5 $s^{-1}$, $Slope_{REHAB/2W}$= 5.2 ±0.3 $s^{-1}$, $Slope_{REHAB/3W}$= 5.2 ±0.3 $s^{-1}$, $Slope_{REHAB/4W}$= 6.0 ±0.2 $s^{-1}$; No significant changes were observed over the 4 weeks period).

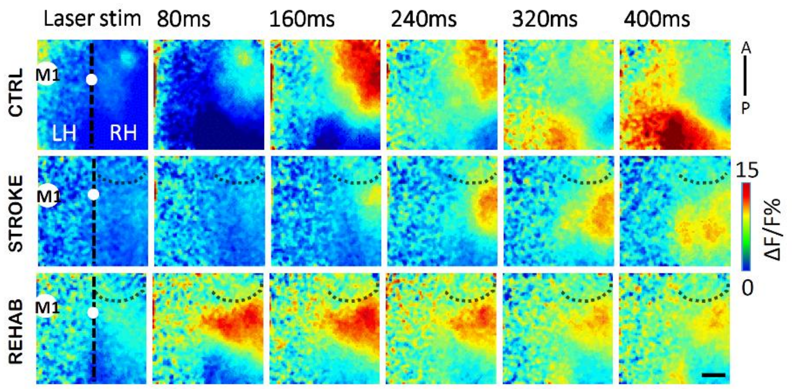


Figure S5. Optogenetic activation of contralateral hemisphere. Related to Figure 5. Representative image sequences from a single animal in each group showing ipsilesional cortical activation as assessed by calcium imaging following 473 nm laser stimulation (1 Hz, 5ms pulse duration) of the ChR2-expressing intact M1. The white spot on the vertical black dotted line (midline) indicates bregma. The dotted curve lines highlight the area injured by stroke. A,P: anterior, posterior. LH, left hemisphere; RH, right hemisphere. Scale bar, 1 mm.

#### Supplemental Experimental Procedures

##### Mice

All procedures involving mice were performed in accordance with the rules of the Italian Minister of Health. Mice were housed in clear plastic cages under a 12 h light/dark cycle and were given *ad libitum* access to water and food. We used two different mouse lines from Jackson Laboratories (Bar Harbor, Maine USA): Tg(Thy1-EGFP)MJrs/J (referred to as GFPM mice) for two-photon imaging experiments and C57BL/6J-Tg(Thy1GCaMP6f)GP5.17Dkim/J (referred to as GCaMP6f mice) for wide-field and optogenetics. Both lines express a genetically-encoded fluorescent indicator controlled by the Thy1 promoter. A subset of GFPM mice imaged for the structural plasticity experiment (dendrites and spines analysis) were used for blood vessels evaluation; a subset of GCaMP6f mice previously used for calcium imaging were analyzed for inter-hemispheric connectivity. Each group contained comparable numbers of male and female mice, and the age of mice was consistent between the groups (4-12 months).

##### Photothrombotic stroke induction

All surgical procedures were performed under Zoletil (50 mg/kg) and xylazine (9 mg/kg) anesthesia, unless otherwise stated. After checking by toe pinching that a deep level of sedation had been reached, the animals were placed into a stereotaxic apparatus (Stoelting, Wheat Lane, Wood Dale, IL 60191). The skin over the skull was cut and the periosteum was removed with a blade. The primary motor cortex (M1) was identified (stereotaxic coordinates +1,75 lateral, -0.5 rostral from bregma). Five minutes after intraperitoneal injection of Rose Bengal (0.2 ml, 10 mg/ml solution in Phosphate Buffer Saline (PBS); Sigma Aldrich, St. Louis, Missouri, USA), white light from an LED lamp (CL 6000 LED, Carl Zeiss Microscopy, Oberkochen, Germany) was focused with a 20X objective (EC Plan Neofluar NA 0.5, Carl Zeiss Microscopy, Oberkochen, Germany) and used to illuminate the M1 for 15 min to induce unilateral stroke in the right hemisphere. Afterwards, the skin over the skull was sutured and the animals were placed in recovery cages until full recovery. The CTRL animals are not subjected to photothrombosis.

##### Optical windows

For the experiments on GCaMP6f mice, we performed a thinned skull preparation on the right hemisphere between bregma and lambda to create an optical window. After applying the local anesthetic lidocaine 2% (20 mg/mL), the skin over the skull and periosteum was removed. The skull over most of the right hemisphere was thinned using a dental drill. A cover glass and an aluminum head- post were attached to the skull using transparent dental cement (Super Bond, C&S). We waited at least 4-5 days after the surgery for the mice to recover before the first imaging session. For the experiments on GFPM mice, we created a square (3x5 $mm^2$) cranial window centered laterally on the right M1 (+1.75 mm from bregma) and extending rostro-caudally from 1 mm posterior to the bregma to lambda. The protocol we followed for cranial window preparation was slightly modified from (Holtmaat et al., 2009) and (Allegra Mascaro et al., 2014). Briefly, we administered anesthetized mice a subcutaneous injection of dexamethasone (0.04 ml per 2 mg/ml). The animals were then placed into a stereotaxic apparatus; after applying the local anesthetic lidocaine 2% (20 mg/mL), the skin over the skull was removed. Using a dental drill (Silfradent, Forlì-Cesena Italia), the border of the area of interest was thinned and the central part of the bone was then gently removed. The exposed brain was covered with a circular cover glass; the optical window was sealed to the skull with a mixture of dental cement and acrylic glue. Finally, an aluminum head-post was attached onto the skull using dental cement (Super Bond, C&S, Sun medical Moriyama City, Shiga, Japan). The surgery was followed by the first imaging session under the two-photon microscope. If the cranial windows were opaque on the second imaging session, the windows were removed and cleaned, and imaging was performed immediately afterwards. After the last imaging session, all animals were perfused with 150 mL of Paraformaldehyde4% (PFA, Aldrich, St. Louis, Missouri, USA).

**Intracortical injections**

We used a dental drill to create a small craniotomy over M1, which was identified by stereotaxic coordinates. Botulinum Neurotoxin E (BoNT/E) injections were performed during the same surgical session in which the photothrombotic lesions were created. We injected 500 nl of BoNT/E (80 nM) divided in 2 separate injections of 250 nl at (i) +0.5 anteroposterior, +1.75 mediolateral and (ii) +0.4 anteroposterior, +1.75 mediolateral at 700 μm cortical depth. For virus injections, we delivered 1 μl of AAV9-CaMKII-ChR2-mCherry ($2.48*10^{13}$ GC/mL) 700-900 μm deep inside the cortex. The skin over the skull was then sutured; the animals were placed in a heated cage (temperature 38°) until they fully recovered.

**Motor training protocol on the M-Platform**

Mice were allowed to become accustomed to the apparatus before the first imaging session so that they became acquainted with the new environment. The animals were trained by means of the M- Platform, which is a robotic system that allows mice to perform a retraction movement of their left forelimb (Spalletti et al., 2014). Briefly, the M-Platform is composed of a linear actuator, a 6-axis load cell, a precision linear slide with an adjustable friction system and a custom-designed handle that is fastened to the left wrist of the mouse. The handle is screwed onto the load cell, which permits a complete transfer of the forces applied by the animal to the sensor during the training session. Each training session was divided into "trials" that were repeated sequentially and consisted of 5 consecutive steps. First, the linear actuator moved the handle forward and extended the mouse left forelimb by 10 mm (full upper extremity extension). Next, the actuator quickly decoupled from the slide and a tone lasting 0.5 s informed the mouse that it should initiate the task. If the animal was able to overcome the static friction (approximately 0.2 N), it voluntarily pulled the handle back by retracting its forelimb (i.e. forelimb flexion back to the starting position). Upon successful completion of the task, a second tone that lasted 1 sec was emitted and the animal was given access to a liquid reward, i.e. 10 µl of sweetened condensed milk, before starting a new cycle.

To detect the movement of the wrist of the animal in the low-light condition of the experiment, an infrared (IR) emitter was placed on the linear slide, and rigidly connected to the load cell and thus to the animal's wrist. Slide displacement was recorded by an IR camera (EXIS WEBCAM #17003, Trust) that was placed perpendicular to the antero-posterior axis of the movement. Position and speed signals were subsequently extracted from the video recordings and synchronized with the force signals recorded by the load cell (sampling frequency = 100 Hz).

All groups performed at least one week (5 sessions) of daily training, starting 26 days after injury for STROKE and TOXIN mice, 5 days after stroke for ROBOT and REHAB group and after the surgery for CTRL animal.

**Optogenetic stimulation and simultaneous recording of GCaMP6f activity**

After the last training session (i.e. 30 days after stroke and at least 2 weeks after the AAV injection in the CTRL group) mice were anesthetized under Zoletil (50 mg/kg) and xylazine (9 mg/kg) and placed into the stereotaxic holder. A small (2x2 $mm^2$) craniotomy was performed over the injected area. After placing the mouse under the wide field fluorescence microscope, we performed repeated laser (473 nm) stimulation (1-2 Hz, pulse duration 3-5 ms, pulse train duration 5 sec, laser power at the focal plane 5 mW) on the left M1, which was localized by mCherry fluorescence. Spurious activation of ChR2 from the green LED (used for GCaMP6f fluorescence excitation) was avoided by blocking half the illumination path with a shutter positioned after the collimator.

**Wide-field fluorescence microscopy**

The custom-made wide-field imaging setup was equipped with two excitation sources for the simultaneous imaging of GCaMP6f fluorescence and light-stimulation of ChR2. For imaging of GCaMP6f fluorescence, a 505 nm LED (M505L3 Thorlabs, New Jersey, United States) light was deflected by a dichroic filter (DC FF 495-DI02 Semrock, Rochester, New York USA) on the objective (2.5x EC Plan Neofluar, NA 0.085, Carl Zeiss Microscopy, Oberkochen, Germany). A 3D motorized platform (M-229 for xy plane, M-126 for z-axis movement; Physik Instrumente, Karlsruhe, Germany) allowed sample displacement. The fluorescence signal was selected by a band pass filter (525/50 Semrock, Rochester, New York USA) and collected on the sensor of a high-speed complementary metal-oxide semiconductor (CMOS) camera (Orca Flash 4.0 Hamamatsu Photonics, NJ, USA).

To perform optogenetic stimulation of ChR2, a 473 nm continuous wavelength (CW) laser (OBIS 473nm LX 75mW, Coherent, Santa Clara, California, United States) was overlaid on the imaging path using a second dichroic beam splitter

(FF484-Fdi01-25x36, Semrock, Rochester, New York USA). The system has a random-access scanning head with two orthogonally-mounted acousto-optical deflectors (DTSXY400, AA Opto-Electronic, Orsay France). A 20X objective (LD Plan Neofluar, 20x/0.4 M27, Carl Zeiss Microscopy, Oberkochen, Germany) was used to demagnify the image onto a 100X100 $pxl^2$ area of the sCMOS camera sensor (OrcaFLASH 4.0, Hamamatsu Photonics, NJ, USA). Images (512x512 pixels, pixel size 9 µm) were acquired at 25 Hz.

**Two-photon fluorescence microscopy**

The custom made apparatus for two-photon microscopy included a mode-locked Ti: Sapphire laser (Chameleon, Coherent Inc.) that supplied the excitation light. The laser beam was scanned in the xy- plane by a galvo system (VM500, GSI Lumonics). An objective lens (XLUM 20X, NA 0.95, WD 2 mm, Olympus) focused the beam onto the specimen. A closed-loop piezoelectric stage (PIFOC ND72Z2LAQ, PhysikInstrumente, Karlsruhe Germany) allowed axial displacements of the objective up to 2 mm with micrometric precision. Finally, the fluorescence signal was collected by a photomultiplier tube (H7710-13, Hamamatsu Photonics). Custom-made software was developed in LabVIEW 2013 (National Instruments).

**Labelling of brain vasculature**

The vasculature was stained using the protocol described by (Tsai et al., 2009), except that we replaced fluorescein (FITC)-conjugated albumin with 0.05% (w/v) tetramethylrhodamine (TRITC)- conjugated albumin (A23016, Thermo Fisher Scientific, Waltham, Massachusetts, USA) in order to avoid spectral overlap between GFP and FITC (Di Giovanna et al., 2018). Under deep anesthesia, mice were transcardially perfused first with 20-30 ml of 0.01 M PBS (pH 7.6) and then with 60 ml of 4% (w/v) paraformaldehyde (PFA) in PBS. This was followed by perfusion with 10 ml of fluorescent gel. After perfusion, a low temperature (ice cold) was maintained to ensure rapid solidification of the gel. After 30 min of cooling, the brain was carefully extracted to avoid damage to pial vessels, and it was incubated overnight in 4% PFA at 4 °C. Dissected cortices were cleared with thiodiethanol (166782 TDE, Sigma Aldrich, St. Louis, Missouri, USA) (Aoyagi et al., 2015; Costantini et al., 2015; Staudt et al., 2007), specifically with serial incubation in 30% and 63% TDE/PBS for 1 h and 3 h, respectively, at room temperature. The cleared cortices were flattened using a quartz coverslip #No1 (UQG Optics).

For the evaluation of endothelial proliferation, REHAB mice received 10 sessions of daily robotic training after BoNT/E injection and stroke induction. REHAB and STROKE animals were transcardially perfused with 4% (w/v) paraformaldehyde (PFA). Brains were post-fixed in PFA and then cryoprotected with 30% (w/v) sucrose for at least 48 hours. 50 µm-thick coronal sections were obtained using a sliding microtome (Leica, Germany). We performed double fluorescence immunohistochemistry (HIC) staining with CD31/Ki67 (CD31, BD Pharmingen, 1:100; Ki67, Abcam, 1:400) to label endothelial cells and verify their proliferation state. We measured the total number of $Ki67^+$ cells in the perilesional area (within 500 µm from the ischemic border), distinguishing those inside blood vessels (labeled by CD31). The results of the analysis are expressed as the fraction of double-labeled cells ($Ki67^+/CD31^+$) over the total number of $Ki67^+$ cells, averaged over the sections (8 sections per animal) for each animal.

**Image analysis**

Two-photon imaging: For the structural plasticity analysis of dendrites and spines of pyramidal neurons, we compared stacks in a vertical mosaic acquired in the rostro-caudal direction. During the last week of training (1st and 4th day), we acquired a mosaic of 100 µm thick stacks (113 x 113 $\mu m^2$) distanced 200 µm from each other along the rostro-caudal axis starting from the borders of the stroke core. We grouped stacks near (<500 µm) and far (from 1000 to 1500 µm) from the core, namely proximal and distal regions, respectively. Dendrite orientation was evaluated on 15 dendrites for each stack (on average) by measuring frame by frame the angle between each structure and the rostro-caudal axis. The stroke core was considered to be at 0°. Z scores were calculated by the Rayleigh Test for circular statistics to evaluate  the angular dispersion of dendrites and blood vessels.

For synaptic plasticity analysis, the fluorescence signal of a spine had to be be at least 1standard deviation higher than the dendritic shaft fluorescence to be included in the analysis. We quantified the plasticity of dendritic spines using two functions: surviving fraction (SF) and turnover ratio (TOR); the SF describes the fraction of persistent structures:

$$SF(t) = N(t2) / N(t1)$$

where N(t1) is the number of spines present during the first imaging session (26 days after injury for STROKE, TOXIN, ROBOT and REHAB mice), while N(t2) indicates those structures that present 4 days after the first imaging session. The TOR evaluates the fraction of newly appeared in the images and disappeared structures:

$$TOR\ (t1, t2) = (Nnew + Ndisappear) / (N(t1) + N\ (t2))$$

where Nnew is the number of structures that are reported for the first time at time t2, Ndisappear is the number of structures that were present at time t1 but which are no longer present at time t2; N (t1) and N (t2) are the total of spines present on t1 and t2, respectively. Unless otherwise stated, data are reported as mean ± SEM.

For vessel orientation analysis, 1 mm deep stacks with 3 µm z-steps were acquired by TPF microscopy. From these acquisitions throughout the entire cortical depth of the right hemisphere, we extracted maximum intensity projections (MIPs; 300 µm thick) over selected sub- stacks, avoiding meningeal vessels in the superficial 100 µm. Four MIPs of proximal regions (within 500 µm from the stroke core), and four MIPs of distal regions (1-1.5 mm from the core) for each mouse were analyzed. We analyzed blood vessels of all sizes, excluding pial vessels. Polar plots of vessel orientation were obtained by measuring frame by frame the angle between each structure and the rostro-caudal axis (n = 30 vessels for each group). For the vessel density analysis, the stacks were first binarized using automatic thresholding with ImageJ. The sum of the pixel count from histograms of the binarized stacks was used as a measure of blood vessel density. Branching points analysis was performed on 3D stacks with the 3D ImageJ suite1 using the workflow for blood vessel segmentation and network analysis developed at the IRB Barcelona (http://adm.irbbarcelona.org/image-j-fiji#TOC-Blood-vessel-segmentation-and-network-analysis). Volumes of 0.53×0.53×0.60 mm were analyzed for both Proximal and Distal regions for each sample (Ollion et al., 2013).

Wide-field calcium imaging: during each experimental session, the mouse's head was restrained and placed on the M-platform under the wide-field microscope. To avoid head movement artefacts, each frame of the fluorescence stack was offline registered by using two reference points (corresponding to bregma and lambda) that were previously marked on the glass window during the surgery procedure.

For each stack (*FluoSt*) a median time series of GCaMP6f fluorescence signal (*mF*) was extracted, where the value of mF at each *i-th* time point corresponded to the median value computed on all the pixels of the *i-th* frame of *FluoSt*. The *mF* was then oversampled and synchronized to the 100 Hz force and position signals.

The *mF* was used to define a GCaMP6f fluorescence signal baseline $F_0$, which was identified by the concomitant absence of fluorescence and force signal deflections. $F_0$was selected within a 2.5 ± 0.7 s interval ($I$) of 62 frames where the fluorescence signal was below 1 standard deviation (STD) of the whole *mF* signal and the corresponding force signal showed a value below of 1 STD of the whole recorded force signal. The fluorescence signal interval was used to reconstruct a 512x512 matrix, i.e. baseline matrix, in which the value of each pixel $B$ of the *{m,n}* coordinates of the matrix was computed as follows:

$$B_{\{m,n\}} = median(p_{\{m,n\},j}) \;\; with\; j = 1, \ldots , N_{\{I\}}$$

where $p_{\{m,n\},j}$ is the value of the pixel $p$ of the *{m,n}* coordinates at the *j-th* frame of the interval $I$ and $N_{\{I\}}$ is the length of Interval $I$. The baseline matrix was then used to normalize all the frames of the fluorescence stack *FluoSt*.

The noise-threshold of 1 STD of the whole recorded force signal was used as a measure of the force peaks exerted by the animal during the retraction task. According to (Spalletti et al., 2013), a force peak is defined as force values that transiently exceed the noise-threshold and result in a movement of the linear slide, as detected by variation of the position signal. To maintain consistency in this analysis, peaks of force that did not result in a movement of the slide were not considered.

The onset of each force peak was used as reference time point to select a sequence of 60 frames (2.4 s, where 0.4 s preceded the force peak) from the *FluoSt*. All sequences were visually checked to exclude possible spurious activation (e.g. early activation or no activation) from the analysis. All the selected sequences (*Seq*s) of the animal *An* on day *d* were compiled, defining a *stack of Seqs,* to compute the *Summed Intensity Projection* for the *An* at *d* (*SIPAn d*). The SIP is a matrix of 512x512 pixels, in which the value *P* of the pixel of the *{m, n}* coordinates is computed as follows:

$$P_{\{m,n\}} = \sum_{k=1}^{Ns} p_{\{m,n\},k}$$

where $p_{\{m,n\},k}$ is the value of the pixel p of the *{m, n}* coordinates at the *k-th* frame of *stack of Seqs,* and $N_s$ is the number of frames of the *stack of Seqs*.

The most active area of the *SIPAn d* was then detected by thresholding the *SIPAn d* with a *median* (*SIPAn d*) + *STD* (*SIPAn d*) threshold value. The threshold *SIPAn d* (*th-SIPAn d*) computed for each week of training on the M-Platform (d=1,…,4 of week W) was superimposed and the common areas, activated at least for 3 daily sessions out of 5 (60%), were labeled as "regions of interest" (ROIs) of the *SIPAn*. We further refined the analysis by dividing the image into two areas, and identifying one anterior ROI [-0.25 - +1.95 mm from bregma (B), AP] and one posterior ROI[+1.95 - + 4.15 mm from B, AP].

We also used the ROIs defined for each individual animal to identify the average ROIs among mice from the same experimental group: *ROIg* with g = "CTRL", "STROKE", "REHAB/1W" and "REHAB/4W" (see Figure S4, panels C and F). Thus, the ROIs from animals of the same group were compiled and further thresholded (60%) to define the *ROIg*. The extent of the *ROIg* was computed as follows:

$$Area_{ROIg} = Area_p \times N_{ROIg}$$

where $Area_p$ corresponds to the area of a single pixel of the image (7.4*10$^{-5}$ mm$^2$) and $N_{ROIg}$ is the number of pixels composing the *ROIg*. Moreover, the centroid of each *ROIg* was identified and its Euclidean distance from bregma was computed (Figure S4, panels D and F).

The ROIs defined for each individual animal were further used to extract the GCaMP6f fluorescence signal corresponding to the activity of those areas. Indeed, from each frame of the FluoS, only pixels belonging to the selected ROI were considered when calculating the representative median value. Thus, a median time series, $F_{ROI}$, was extracted from the whole FluoS and was representative of the ROI. The fluorescence signal was normalized ($\frac{\Delta F_{ROI}}{F_0}$ 100%) and low-pass filtered to clean the signal from the detected breathing artefacts (Chebyshev filter with cutting frequency = 9 Hz). The previously detected force peaks were then used to select the GCaMP6f fluorescence peaks from the $\frac{\Delta F_{ROI}}{F_0}$ signal.

A time window that lasted 4 seconds, i.e. *wnd*, and was centered at the onset of the force peak, was used to delimit a part of the $\frac{\Delta F_{ROI}}{F_0}$ signal , i.e. $F_{wnd}$, to identify the corresponding fluorescence peak. A fluorescence peak was defined as the part of $F_{wnd}$ that overcame the value of *median+3STD* calculated for the whole signal $\frac{\Delta F_{ROI}}{F_0}$.The onset of the peak was detected as:

$$t_{\{peak\ onset\}} = t_{\left\{min\left(\frac{dF_{wnd}}{dt}\right)\right\}}$$

calculated for the $[t_{st}\ t_{max}]$ time interval, where $t_{st}$ and $t_{max}$ are the time points corresponding to the start of the *wnd* and the maximum of $F_{wnd}$ , respectively.

From the fluorescence and force peaks, different parameters were computed as follows:

- the maximum of the peak (*Peak amplitude*).
- the *Slope* of the peak was defined as follows:

$$Slope_{\{peak\}} = max\left(\frac{dS}{dt}\right)$$

between the $t_{\{S\ peak\ onset\}}$and $t_{S\ max}$ where *S* = *fluo* or *force* signal.

- the time delay between the occurrence of the maximum of the fluorescence (*ΔT*).

The first movement time point was defined as the first variation of the position signal ($\frac{dx}{dt} > 0$) detected along the *wnd* interval.

Optogenetically-induced calcium activity: On every stack, we analyzed the calcium activity averaged over a round ROI (2.3 mm of diameter) centred on the peri-infarct area. Changing in the variation of fluorescence signal (ΔF/F) triggered by light irradiation of the contralateral hemisphere that were below 1% were excluded from the analysis. A calcium response was considered optogenetically-triggered if it started within 120 ms of optogenetic stimulation. The activation delay refers to the average (± Standard Error of the Mean, SEM) delay of the calcium peak with respect to the onset of laser stimulation. The success rate reports the number of times the laser stimulation successfully triggered contralateral activation over the total number of stimulation trials.

SI Bibliography


Allegra Mascaro, A.L., Sacconi, L., and Pavone, F.S. (2014). Laser nanosurgery of cerebellar axons in vivo. Journal of visualized experiments : JoVE, e51371.

Aoyagi, Y., Kawakami, R., Osanai, H., Hibi, T., and Nemoto, T. (2015). A rapid optical clearing protocol using 2,2'-thiodiethanol for microscopic observation of fixed mouse brain. PloS one *10*, e0116280.

Costantini, I., Ghobril, J.P., Di Giovanna, A.P., Allegra Mascaro, A.L., Silvestri, L., Mullenbroich, M.C., Onofri, L., Conti, V., Vanzi, F., Sacconi, L.*, et al.* (2015). A versatile clearing agent for multi-modal brain imaging. Scientific reports *5*, 9808.

Di Giovanna, A.P., Tibo, A., Silvestri, L., Müllenbroich, M.C., Costantini, I., Allegra Mascaro, A.L., Sacconi, L., Frasconi, P., and Pavone, F.S. (2018). Whole-Brain Vasculature Reconstruction at the Single Capillary Level. Scientific Reports *8*, 12573.

Holtmaat, A., Bonhoeffer, T., Chow, D.K., Chuckowree, J., De Paola, V., Hofer, S.B., Hubener, M., Keck, T., Knott, G., Lee, W.C.*, et al.* (2009). Long-term, high-resolution imaging in the mouse neocortex through a chronic cranial window. Nature protocols *4*, 1128-1144.

Ollion, J., Cochennec, J., Loll, F., Escude, C., and Boudier, T. (2013). TANGO: a generic tool for high-throughput 3D image analysis for studying nuclear organization. Bioinformatics *29*, 1840-1841.

Spalletti, C., Lai, S., Mainardi, M., Panarese, A., Ghionzoli, A., Alia, C., Gianfranceschi, L., Chisari, C., Micera, S., and Caleo, M. (2014). A robotic system for quantitative assessment and poststroke training of forelimb retraction in mice. Neurorehabilitation and neural repair *28*, 188-196.

Spalletti, M., Comanducci, A., Vagaggini, A., Bucciardini, L., Grippo, A., and Amantini, A. (2013). Efficacy of lacosamide on seizures and myoclonus in a patient with epilepsia partialis continua. Epileptic disorders : international epilepsy journal with videotape *15*, 193-196.

Staudt, T., Lang, M.C., Medda, R., Engelhardt, J., and Hell, S.W. (2007). 2,2'-thiodiethanol: a new water soluble mounting medium for high resolution optical microscopy. Microscopy research and technique *70*, 1-9.

Tsai, P.S., Kaufhold, J.P., Blinder, P., Friedman, B., Drew, P.J., Karten, H.J., Lyden, P.D., and Kleinfeld, D. (2009). Correlations of neuronal and microvascular densities in murine cortex revealed by direct counting and colocalization of nuclei and vessels. The Journal of neuroscience : the official journal of the Society for Neuroscience *29*, 14553-14570.
